# Climate of origin constrains phenological tracking of temperature in a California endemic oak (*Quercus lobata*)

**DOI:** 10.64898/2026.09.26.754716

**Authors:** Alexander R. B. Goetz, Victoria L. Sork, Courtney Canning, Christopher T. Ivey, Eli Carlisle, Eric Wada, Jessica W. Wright

## Abstract

**Premise of the study:** Phenotypic plasticity is advantageous for many plants that experience environmental fluctuations, but strong selection pressures can establish limits to plasticity. In deciduous trees, the ability to time leaf elongation in response to spring temperatures can extend the length of the growing season for a given year’s climate. However, emergence timing may have undergone strong selection by factors like cold temperatures, limiting the capacity of locally adapted populations to adjust via plasticity to new conditions.

**Methods:** Using a nine-year dataset of 659 families of *Quercus lobata* in two common gardens, we tested how climate of origin and annual temperatures at the gardens affected variation in phenology. We then examined whether leaf emergence timing was under selection in the gardens by testing its effects on growth. Finally, we tested whether among-year plasticity in leaf emergence timing conferred a growth benefit.

**Key results:** Year-to-year environmental variation explained more phenological variation than genetic differences associated with climate of origin, yet trees from cooler climates were more constrained in their ability to track temperature than trees from warmer climates. We found evidence of weak stabilizing and directional selection on phenology, suggesting that strong selection in favor of earlier phenology is unlikely. Finally, we found no evidence that selection has favored increased plasticity in *Q. lobata*.

**Conclusions:** Phenological plasticity remains a key driver of plants’ response to changing temperatures, but it is constrained by natural selection. As temperatures rise, these processes may disproportionately limit the ability of trees adapted to cooler climates to shift their phenology to match spring temperatures.

## INTRODUCTION

Phenotypic plasticity allows many plant species to rapidly respond to fluctuating environmental conditions, and the limits of that plasticity may predict the degree to which their fitness is affected by environmental change. Plastic responses to the environment, such as shifts in phenology, may be especially important for species with long generation times (Alpert and Simms, 2002; Gimeno et al., 2008; Nicotra et al., 2010) because these species cannot rapidly respond to environmental change through dispersal or natural selection (Jump and Peñuelas, 2005; Aitken et al., 2008; Sork et al., 2013). Phenological plasticity may be beneficial if it allows plants to “track” environmental variation by adjusting the timing of phenological events in response to changing environmental cues (Anderson et al., 2012; Cleland et al., 2012; Wolkovich and Donahue, 2021). In contrast, if plants do not adjust their phenology in response to warming temperatures, they may not experience the full growing season that could result in increased annual growth (Larcher, 2003; Augspurger, 2008; Inouye et al., 2019). Thus, if warming temperatures reduce the frequency and severity of early freeze events, we may expect relaxation of the selective pressures that constrain the tracking of temperature cues, potentially increasing the importance of plasticity in determining phenology (McDonough MacKenzie et al., 2018; Gao et al., 2023; Gao et al., 2025). Alternatively, tracking may still be constrained by past selection against early leaf emergence (Alpert and Simms, 2002; Valladares et al., 2007; Dantec et al., 2015; Körner et al., 2016) or by the effects of photoperiod cues (Basler and Körner, 2012; Zohner and Renner, 2014; Flynn and Wolkovich, 2018). Determining what constrains phenological tracking can reveal the limits of plasticity in trees and how those limits may affect how their capacity to keep pace with rapid changes.

One potential constraint on phenological tracking is local adaptation to climate, which can generate genetic differences in the phenological responses to environmental change. Spring leaf phenology reflects a fundamental trade-off wherein earlier leaf elongation extends the growing season, which is expected to increase annual growth, but also increases the risk of stress or damage from cold temperatures, herbivory, or disease (Inouye et al., 2019). Cold temperatures impose an especially strong selective pressure that constrains the onset of spring leaf elongation (Dantec et al., 2015; Körner et al., 2016; Lenz et al., 2016). In addition, biotic consequences of early phenology, such as increased risk of disease or insect herbivory, can cause significant damage to trees (Marquis and Whelan, 1994; Dantec et al., 2015; Renner and Zohner, 2018). Therefore, the costs of leafing out too early, which may include mortality or severe tissue damage, are expected to exceed the costs of leafing out too late, which may reduce growth (Inouye et al., 2019). These asymmetric costs can generate strong selection on phenology. Accordingly, many species show substantial genetic variation in phenology (Ducousso et al., 1996; Jensen and Hansen, 2008; Morin et al., 2010) associated with local adaptation to climate, with populations originating from cooler climates exhibiting later phenology than warm-climate populations (Rehfeldt, 1989; Farmer, 1993; Chuine, 2010).

For plants to track changing climate through altered phenology, they must respond plastically to the cues that induce leaf development. Initial budburst reflects the initiation of spring leaf development, and buds are most vulnerable to damage at this stage (Körner et al., 2016; Hänninen et al., 2019). Complete emergence of leaves from the bud (“leaf elongation”) represents a later stage of development associated with an increase in photosynthetic activity (Lechowicz, 1984). The timing of budburst and leaf elongation is influenced by chilling, warming, and day length (photoperiod) (Flynn and Wolkovich, 2018; Wolkovich et al., 2022), with temperature often being a predominant factor (Laube et al., 2014; Nanninga et al., 2017; Ettinger et al., 2020; Guo et al., 2020). Although photoperiod can also influence spring leaf phenology (Basler and Körner, 2012; Flynn and Wolkovich, 2018), responsiveness to temperature is particularly important for tracking climate change because, unlike temperature, photoperiod does not change with climate (Zohner and Renner, 2014). Thus, plastic responses to temperature may allow plants to adjust the timing of leaf development and growing season length as climates warm (Cleland et al., 2012; Wang et al., 2023).

Given that both adaptation to historical climate conditions and plasticity shape spring leaf phenology, populations may differ in their capacity to track changing temperatures. Phenology is especially sensitive to interannual environmental variation (Morin et al., 2010; McDonough MacKenzie et al., 2018) and short-term extremes (Carter et al., 2017; Ladwig et al., 2019). In multiple species, phenology is more variable among individuals in warmer years (Gao et al., 2023; Gao et al., 2025). However, climate-associated genetic variation may constrain this plastic response, causing populations to differ in their capacity to track climate change (Zohner and Renner, 2014; Ramos-Muñoz et al., 2025). In many species, populations originating from warmer environments have shown greater plasticity in traits that associated with responses to rising temperatures (Lloret et al., 2011; Nicotra et al., 2015), including phenological tracking (Morin et al., 2009). Such differences in plasticity may persist if historical selection has imposed genetic constraints on phenology or if plasticity itself carries fitness costs (DeWitt et al., 1998); Valladares et al. (2002). Conversely, warming could relax selection against early leaf emergence, potentially reducing these constraints. Determining whether contemporary selection on phenology maintains these constraints is therefore critical for understanding the capacity of populations to track ongoing climate change (Cleland et al., 2012; Inouye et al., 2019; Iler et al., 2021).

The overarching goal of this study was to determine whether climate of origin constrains phenological tracking of temperature and whether contemporary selection maintains these constraints. While previous studies have found evidence of genetic differentiation in phenological plasticity among local populations, including in oaks (Vitasse et al., 2009; Morin et al., 2010; Vitasse et al., 2013; Knott et al., 2023), tests across broad climatic gradients, many families, and multiple years remain limited, constraining our ability to predict species-wide phenological responses to climate change (Tang et al., 2016; Hänninen et al., 2019). To assess genetic and plastic contributions to leaf phenology, we measured spring budburst and leaf elongation in progeny from 659 families of *Quercus lobata* Née (valley oak) growing in two common gardens between 2015 and 2026. The two common gardens differed in temperature, among other variables (Delfino Mix et al., 2015), and only one garden experienced spring freezing temperatures. Families sampled across the species range represented a broad gradient of source climates, while repeated measurements across years exposed trees to substantial interannual temperature variation and allowed us to evaluate phenology beyond the seedling stage. *Quercus lobata* is well suited to this test because the species occupies a wide climatic niche, including substantial variation in winter and spring minimum temperatures (Pavlik et al., 1991; Goetz et al., 2026), shows strong signals of genetic adaptation to local environments (Sork et al., 2010b; Gugger et al., 2016; Gugger et al., 2021), and displays substantial levels of plasticity and genetic variation in several traits (MacDonald, 2017; Browne et al., 2019; Ramirez et al., 2020; Browne et al., 2021; Ochoa, 2024), including phenology (Koenig et al., 2021; Wright et al., 2021).

To address our central goal, we utilized a long-term common garden study of a widespread California oak to ask three related questions: (1) Does climate of origin predict genetic differences in spring phenology? (2) Does climate of origin constrain the capacity of trees to track interannual temperature variation through phenotypic plasticity? (3) Does contemporary selection on phenology or phenological plasticity help explain maintenance of these constraints? By analyzing phenology across years by examining families spanning the species’ climatic range and growing in two common gardens, we test both the magnitude and limits of phenological tracking in response to temperature variation. We find that interannual variation exerts a greater influence on *Q. lobata* leaf phenology than genetic differences associated with climate of origin, but that this plasticity is constrained: Trees originating from cooler climates show reduced capacity to track temperature variation.

## MATERIALS AND METHODS

### Study species

*Quercus lobata* Née is a winter-deciduous tree oak endemic to woodlands, savannas, and riparian areas of the California Floristic Province (Pavlik et al., 1991). As a foundational species in the California oak savanna and riparian ecosystems, it supports extraordinary biodiversity (Allen-Diaz et al., 1999). The species is found across wide abiotic gradients (0-1700m above sea level; Pavlik et al. 1991). *Quercus lobata* has experienced extensive habitat reduction across its range, primarily due to land use change (Tyler et al., 2006; Whipple et al., 2011) and natural populations have shown limited recruitment (Tyler et al., 2006). In addition, populations appear maladapted to current climate conditions, possibly due to an adaptational lag to cooler historical temperatures (Browne et al., 2019; Goetz et al., 2026), which raises concerns about the vulnerability of *Q. lobata* to warmer climates. Landscape genomic analysis has identified regions where *Q. lobata* will be even more vulnerable to climate warming (Buck et al. *in prep*).

### *Quercus lobata* common garden study

The common garden study of *Q. lobata* was established by VL Sork and JW Wright in 2012 (described in Delfino Mix et al. 2015). Acorns were collected from 674 mature *Q. lobata* individuals across the species range and planted into two common gardens administered by the USDA Forest Service (“Chico garden” at the Chico Seed Orchard, Chico, CA, 70m above sea level; “Placerville garden” at the Institute of Forest Genetics, Placerville, CA, 822m above sea level; Figure 1). The Placerville garden typically experiences colder temperatures from December to May than the Chico garden (Figure S1). In 2015, two-year-old seedlings were planted into 2-ha fields following one year of growth in a greenhouse at IFG and a second year of growth in lath houses at the respective garden sites. In 2021, roughly half of the trees were deliberately thinned as per the original study plan to reduce canopy competition. As of 2025, a total of 3,673 surviving half-sib progeny of 659 families were present across both gardens.

**Figure 1:**
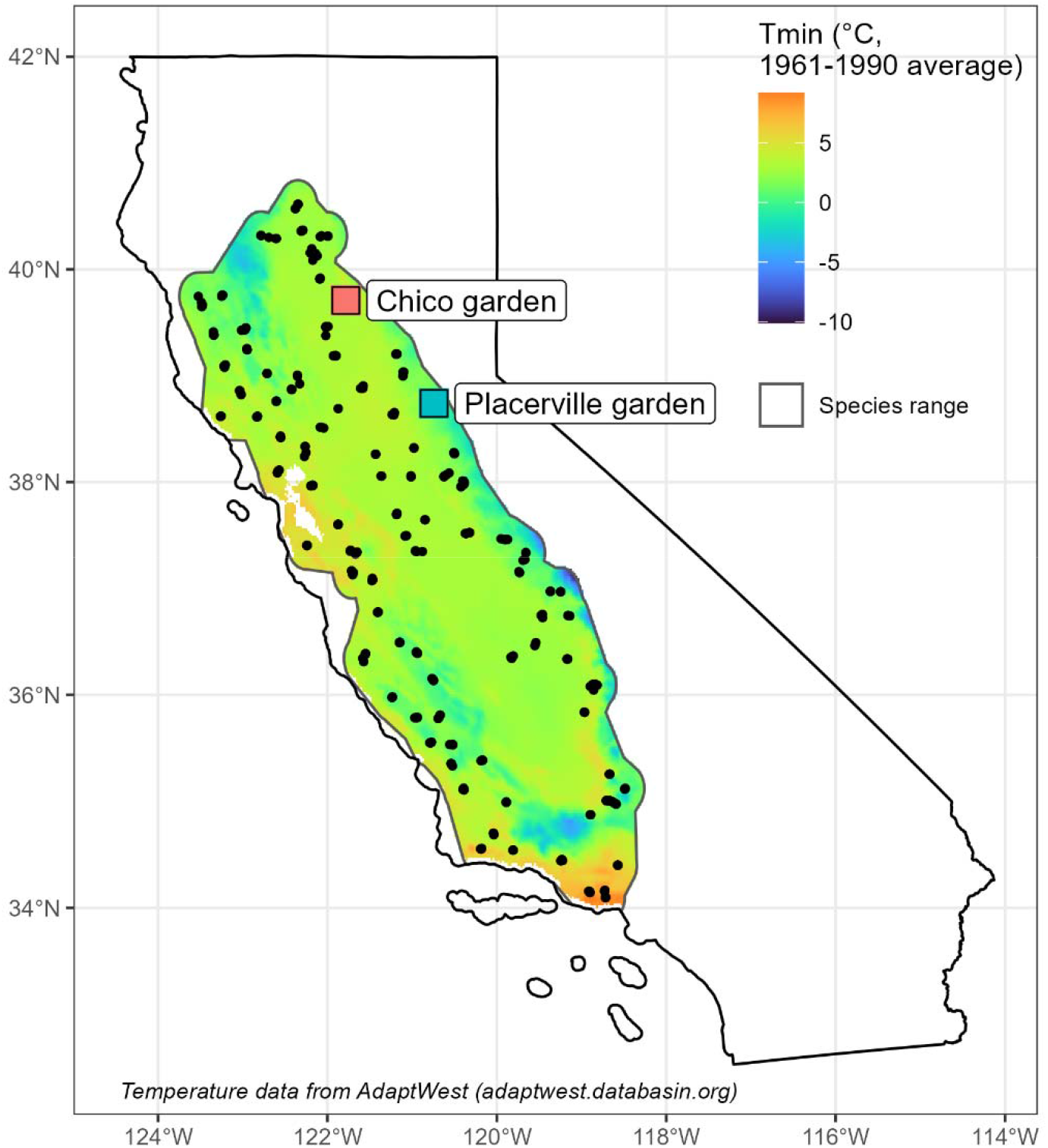
Localities (black circles, n = 95) of the midpoints of clusters of 3-5 l *Q. lobata* trees used as seed sources for 659 families planted into two common gardens (labeled squares). Outline represents the theorized current species range based on the distribution of living trees (Davis *et al*. 1998). Labeled squares represent the two common garden locations (red: Chico Seed Orchard, Chico, CA; blue: Institute of Forest Genetics, Placerville, CA). Background color represents recent-past minimum annual temperature averaged from 1961-1990 (AdaptWest, 2022).

**Figure 2:**
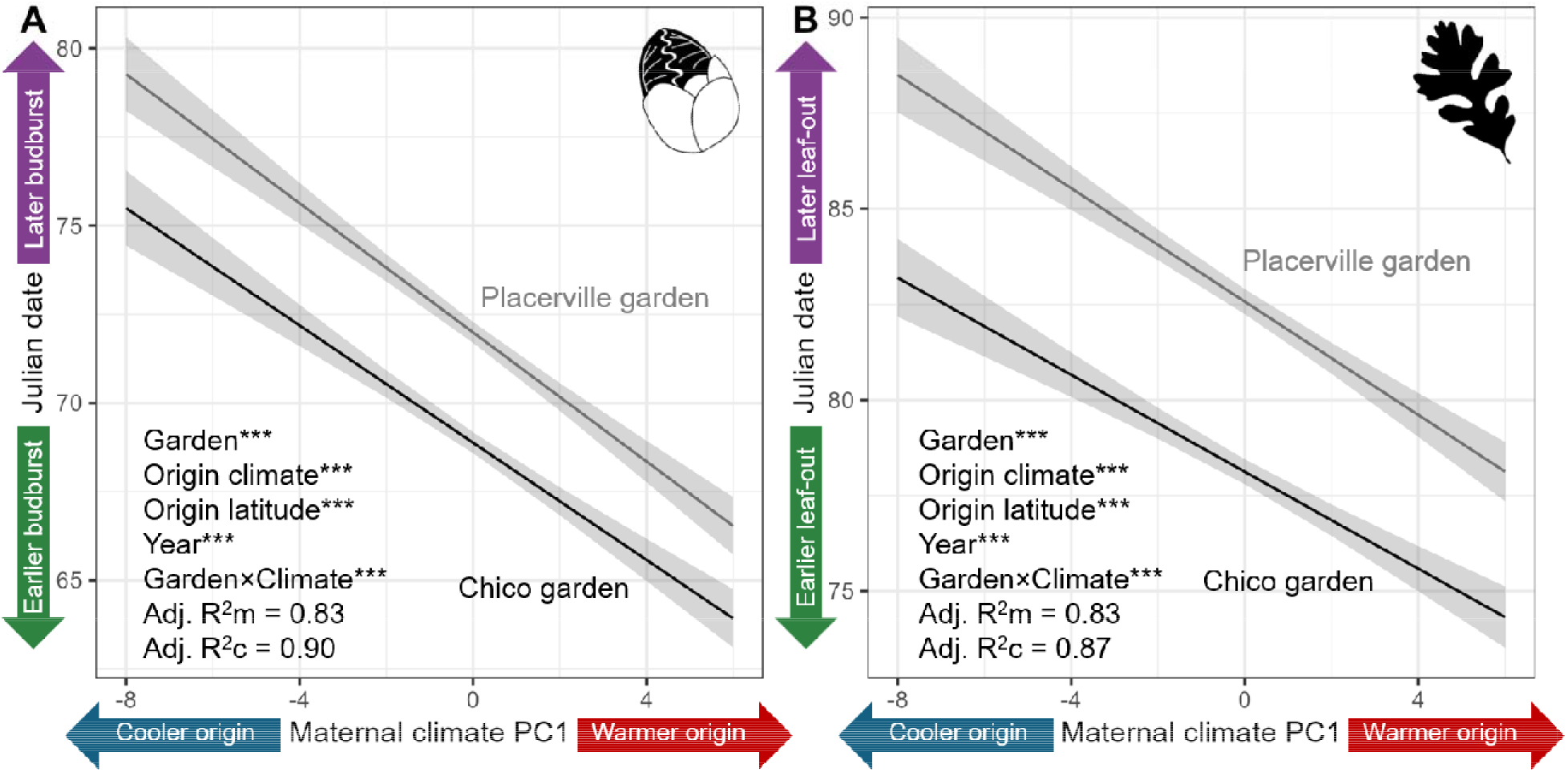
Trees from warmer, drier origins have earlier date of budburst and leaf elongation in the common gardens than trees from cooler, wetter origins. Lines represent the marginal effect of the interaction term between maternal origin climate and garden (black: Chico, grey: Placerville) on dates of phenological events, from a repeated-measures mixed model that also accounted for latitude of origin and study year. In each model, there were significant effects of garden, study year, climate of origin, and site*climate of origin interaction on dates of phenological events (P < 0.01). Shaded intervals are 95% confidence intervals. Model summaries are provided in Table S5. **A**: Date of first budburst. **B**: Date of first leaf elongation.

The quantitative genetic design of our study focuses on differences among families, with individual maternal trees and their site-specific climate environments as the scope of inference in analyses. Within localities, we sampled families far enough apart (Delfino Mix et al., 2015) so that they are genetically unrelated (Dutech et al., 2005) and we scaled climate variables to the tree level rather than locality level. Therefore, we test clinal effects of climate on phenotypes in the gardens at an individual tree scale instead of treating localities as distinct genetic subpopulations.

### Data collection

#### Phenology of leaf elongation

Between 2015 and 2026, field crews visually surveyed the stages of spring leaf elongation of all trees in the Placerville garden during 10 seasons (2015, 2016, 2017, 2018, 2019, 2020, 2021, 2024, 2025, 2026) and in the Chico garden during 7 seasons (2015, 2016, 2018, 2019, 2024, 2025, 2026; financial constraints and COVID-19 restrictions prevented collections during some years). The stages of bud burst through leaf elongation were categorized as described by Derory et al. (2006): 0 = fully closed buds, 1 = bud swelling (visible green tissue) 2 = bud opening; 3 = leaf venation visible within bud; 4 = at least one leaf fully out of bud, 5 = fully elongated leaves (visible internodes). Surveys were conducted once per week February and May of each year from 2015-2021, and twice per week from 2024-2026. To compare date of budburst and leaf elongation among individual trees and families, we calculated the date of first evidence of budburst (typically, but not always, stage 1) and leaf elongation (stage 5) as the number of days since January 1st. To allow us to test the effect of selection on phenology, we also calculated each tree’s average date of budburst and leaf elongation across all study years (excluding years with incomplete data). Refer to Table S1 for details on sample sizes, survey date ranges, and observers in each year.

#### Maternal origin climate

To investigate whether local adaptation to the origin climate affects phenology in the common gardens, we used 10 climate variables to characterize the origin climate of the maternal seed source: Maximum summer temperature, minimum winter temperature, maximum, minimum, and average annual temperatures, climatic water deficit, temperature seasonality, precipitation seasonality, precipitation of warmest quarter, and precipitation of coldest quarter. These variables were previously shown by (Browne et al., 2019) to accurately characterize variation in local climate conditions of *Q. lobata* populations. We extracted 30-year averages of each climate variable for the point locations of each maternal tree from the Basin Characterization Model (BCM; Flint et al., 2013) at 270m pixel resolution. We used temperature averages from the 1951-1980 period because they are closer to the conditions in which the maternal trees (many of which are estimated to be over 100 years old) would have germinated (Browne et al., 2019; Goetz et al., 2026).

#### Garden climate

To test whether interannual climate variation within and between gardens affects phenology, we obtained daily maximum and minimum temperatures, as well as daily precipitation values, at the garden sites during all study years. Data were obtained from PRISM (prism.oregonstate.edu) using the Explorer tool, which provides time series data for point locations extracted from 800m rasters with interpolation. To explicitly compare date of budburst and leaf elongation with climatic conditions at relevant time points, we calculated averages of minimum and maximum monthly temperature as well as total precipitation for winter (December-February) and spring (March-May). We verified that the PRISM data accurately reflected conditions at the garden plots by fitting a linear regression between PRISM values and values collected using a series of iButton (iButtonLink Technology, Whitewater, WI, USA) temperature sensors at the gardens between 2015 and 2019 (Figure S2).

#### Relative growth rates

To test for selective pressure on phenology using a component of fitness, we calculated relative growth rates (RGR) based on tree size. Tree size has been measured on all trees in both gardens annually since 2013. Heights of all trees were measured between 2013 and 2017; starting in 2018, some trees were too tall to measure, so diameter at breast height (1.4m; DBH) was measured instead for those trees. Heights of all trees were measured in 2024 and 2025 at the Placerville Garden. Basal diameter was measured on all trees from 2013-2015 and on a haphazard subset of trees in 2018. Thus, to allow for comparisons of tree size change across years, we used predictive models of allometric relationships between basal diameter, DBH, and height to calculate a standardized size metric, height’, from which we calculated single-season and cumulative relative growth rates (refer to Goetz et al., 2026 for more details).

### Statistical analysis

#### Software

All analyses were carried out using R version 4.5.0 (R Core Team, 2025) with RStudio version 2025.09.1 (Posit Posit team, 2025). All linear mixed models were fitted using function “lmer” in R package “lmerTest” version 3.2-1 (Kuznetsova et al., 2017), which imports package “lme4” version 2.0-1 (Bates, 2010; Bates et al., 2015). We used function “r.squaredGLMM” in package “MuMIn” version 1.48.19 (Bartoń, 2026) to calculate marginal and conditional R^2^ values for all linear mixed models. We used function “ggpredict” in package “ggeffects” version 2.3.2 (Lüdecke, 2018) to calculate all model predictions. Figures were created using packages “ggplot2” version 4.0.3 (Wickham, 2016) and “sjPlot” version 2.9.0 (Lüdecke, 2024).

#### Principal components analysis of origin climate

To summarize the variation in origin climate for use in analysis, we calculated principal component vectors (PCs) after (Browne et al., 2019). The principal component analysis was calculated based on 10 climate variables: Maximum summer temperature, minimum winter temperature, maximum, minimum, and average annual temperatures, climatic water deficit, temperature seasonality, precipitation seasonality, precipitation of warmest quarter, and precipitation of coldest quarter, all derived from the Basin Characterization Model (Flint et al., 2013). The first PC vector (40.8% of variance explained) represented the spectrum between hotter, drier sites and cooler, wetter sites, while the second PC vector (23.3% of variance explained) represented seasonal differences in temperature and precipitation (higher values indicate temperature-based seasonality, lower values indicate precipitation-based seasonality). Refer to Figure S3b for PCA biplot and Table S2 for variable loadings.

#### Genetic patterns in date of budburst and leaf elongation

To determine whether date of budburst and leaf elongation are genetically determined, we utilized our quantitative genetic design to test whether each variable differed among families and partitioned the variance components to estimate quantitative trait values (QST; Falconer and Mackay, 1983). First, we modeled the difference in date of budburst and leaf elongation among families in each garden, using a mixed model with family as a random variable. To primarily test phenology differences not caused by temperature fluctuations, we used each tree’s average dates of budburst and leaf out over the study period as the dependent variables. To account for differences due to maternal effects, we included each tree’s initial height prior to outplanting. Because the Chico garden consists of three separate sections that vary environmentally, we included section as an additional fixed effect in the model for the Chico garden. We then partitioned variance components using function “VarCorr” from package “lme4” and calculated *Q*_ST_ using the following formula (Falconer and Mackay, 2009):

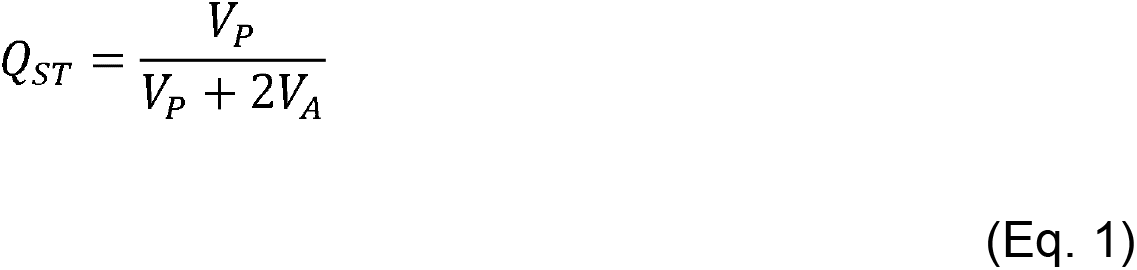

Where V_P_ is total estimated phenotypic variance and V_A_ is estimated additive genetic variance (phenotypic variance explained by differences among families).

To determine whether differences were significant, we conducted restricted likelihood ratio tests (RLRT; (Morrell, 1998) on the mixed models used to calculate *Q*_ST_ values. RLRTs were conducted using function “exactRLRT” from package “RLRsim” version 3.1-9 (Scheipl et al., 2008). The restricted likelihood ratio test is a robust method to test for significant differences among levels of a random factor (Morrell, 1998).

To test for genotypic, environmental, and genotype-by-environment effects on date of budburst and leaf elongation among years, we conducted repeated-measures mixed models on the three-way interaction between family, garden, and study year. To explicitly test reaction norms among the two sites, we also modeled the effect of the garden-by-family interaction on trees’ average dates of budburst and leaf elongation across all study years. We fit models using function “lmer” in R package “lmerTest” and tested significance using function “anova” in R package “stats.”

To compare date of budburst and leaf elongation at the family level while accounting for sampling artifacts, we calculated best linear unbiased predictors (BLUPs) for each family in each site in each year. To calculate BLUPs, we fit linear mixed models on dates of first budburst and first leaf elongation separately by site. Both models included the random effects of family and observer. Because the Chico garden consists of three separate sections that vary environmentally, we included section as a random variable in the model for the Chico garden.

#### Effects of maternal climate on date of budburst and leaf elongation

To test whether trees from warmer sites consistently burst bud and leaf out earlier, we fit repeated-measures linear mixed models (Gezan and Carvalho, 2018) on the relationship between source climate and BLUPs of date of budburst or leaf elongation. We quantified source climate as the first principal component vector (PC1) of 10 climate variables (“Principal components analysis” above). Higher values of PC1 indicate hotter, drier origin sites, while lower values indicate cooler, wetter origin sites. We fit models separately for each garden because large differences in timing of spring warming between gardens caused discontinuity if data were pooled. In both models, we included garden as an interaction term with origin climate to test whether the effects of origin climate differed across growing environments. We also included origin latitude as a covariate because it was correlated with PC1 (Figure S3) and to test for photoperiod effects on climate adaptation.

#### Effects of interannual temperature variation on date of budburst and leaf elongation

To test whether trees generally have earlier phenology in warmer years at the gardens, we fit linear mixed-effects models relating yearly garden temperatures to budburst and leaf elongation dates. First, to determine which abiotic cue (maximum temperature, minimum temperature, or precipitation) best explained yearly variation in date of budburst and leaf elongation, we ran univariate linear regressions on the relationship between each phenology variable and each abiotic variable and compared the resulting models based on Akaike’s Information Criterion (AIC, Akaike, 2025).

For the budburst models, we used the minimum temperature averaged between December and February of each year, and for the leaf elongation models we used minimum temperature averaged between March and May of each year. The models used pooled data from both common gardens. As before, we included maternal family as a random effect.

#### Effects of source temperature on response to interannual temperature variation

To test whether trees responded differently to annual garden temperatures based on their maternal source temperatures, we fit generalized additive models (GAMs; Hastie and Tibshirani 1986) relating the interaction between garden temperature and source temperature to date of budburst and leaf elongation. To model the interaction term in two dimensions, we used a tensor spline (Wood, 2017). As before, we included maternal family as a random effect and pooled data from both common gardens. Models were fitted using function “bam” in R package “mgcv” version 1.9-4 (Wood, 2011, 2017). We used function “gam.check” from “mgcv” to evaluate model fits and used function “fitted_values” from R package “gratia” version 0.11.2 (Simpson, 2024) to obtain model fits for visualization.

#### Contemporary selection on leaf phenology

To test whether date of budburst and leaf elongation are under selection in the garden environments, we fit linear models relating relative growth rates (RGR) to date of budburst and leaf elongation. First, we tested whether families’ average date of budburst and leaf elongation across years affected their cumulative RGR between 2014 and 2025 separately for the two gardens separately. To identify whether selection at the two gardens was directional or stabilizing, we tested the significance of the phenology variables as both linear and quadratic terms and compared their significance and R^2^ values in the models. Each model accounted for maternal family (random effect), summer maximum temperature at the maternal origin site (because trees from warmer sites have been previously shown to have higher RGR than those from cooler sites; Browne et al., 2019; Goetz et al., 2026), and initial height at the start of the study period (t = 2014 in the cumulative models, t = the prior growing season in the annual models).

#### Costs or benefits of plasticity

To test whether plasticity imposes costs or benefits to growth, we used linear models to test the effect of coefficient of variation (CV) of each family’s dates of budburst and leaf elongation with their cumulative RGR. These coefficients of variation measure the degree to which an individual tree’s phenology is plastic over time. A negative relationship between CV and RGR would indicate that higher plasticity (greater variation in the tree’s budburst or leaf elongation date among years) imposes a cost to growth, while a positive relationship would indicate that higher plasticity provides a growth benefit. Finally, no relationship would indicate that individual plasticity does not affect cumulative growth. We calculated CV for each tree based on the variance and mean of their date of budburst and leaf elongation across years, then calculated average CV by family. We also tested whether origin climate affects CV using linear regression.

## RESULTS

### Date of budburst and leaf elongation are genetically determined

Date of budburst and leaf elongation significantly differed among families in each garden, with *Q*_ST_ values ranging from 0.24 to 0.28 (Table 1). In addition, date of budburst and leaf elongation showed significant differences across years and a significant interaction between family and year of study (Figure S4; Table S3), indicating that families showed different phenological patterns across warmer and cooler years. We observed evidence of phenotypic plasticity between gardens, but no significant interaction between family and garden (Figure S4; Table S4). In sum, we find evidence for a genetic basis to their phenology in *Q. lobata*, consistent with previous findings by Wright et al. (2021). These results raise the question of whether selection associated with trees’ environments of origin has shaped phenology.

**Table 1:** Significant differences in date of budburst and leaf elongation among 659 families of *Quercus lobata* trees growing in two common gardens. Variance components, derived *Q*_ST_ values, and results of a restricted likelihood ratio test on linear mixed models testing the random effect of family on budburst and leaf elongation dates (average per individual) measured among *Q. lobata* half-sib families in a common garden. The fixed effect of tree height prior to outplanting was also included in the model to account for maternal effects on trees’ early growth. At the Chico garden, the additional fixed effect of planting section was included to account for known environmental differences within the plot. V_A_: Additive genetic variance (4*V_Family_). V_P_: Total phenotypic variance (V_Family_ + V_Planting block_ + V_Residual_). Q_ST_: V_P_ / (V_P_ + 2*V_A_) (Falconer and Mackay, 2009). RLR: Restricted likelihood ratio (Morrell, 1998). P: P-value from restricted likelihood ratio test of variance among families, based on 10,000 simulated values.

| Garden | Variable | $V_A$ | $V_P$ | $Q_{ST}$ | RLR | P |
| --- | --- | --- | --- | --- | --- | --- |
| Placerville | Date of first budburst | 50.9 | 31.5 | 0.24 | 212.8 | < 0.001*** |
| Placerville | Date of full leaf elongation | 58.3 | 39.1 | 0.25 | 189.2 | < 0.001*** |
| Chico | Date of first budburst | 49.4 | 35.6 | 0.27 | 189.4 | < 0.001*** |
| Chico | Date of full leaf elongation | 31.8 | 24.9 | 0.28 | 160.5 | < 0.001*** |

### Trees from warmer, drier origins burst bud and leaf out earlier

Budburst and leaf elongation dates were strongly correlated with climate of origin in all study years and at both gardens (Figure 3). Trees from warmer, drier environments consistently burst bud earlier than trees from cooler, wetter environments. We found significant but small interaction effects of garden (budburst: P < 0.01; leaf elongation: P = 0.02; Table S5), with a slightly stronger effect of origin climate on phenology at the Placerville garden than the Chico garden. We also found a significant but small effect of origin latitude on date of budburst and leaf elongation (P < 0.001 for both budburst and leaf elongation; Table S5); trees from the southernmost extent of the range burst bud roughly 2.5 days later than trees from the northernmost extent of the range (Figure S5). Finally, trees at the Placerville garden consistently exhibited later phenology than trees at the Chico garden (budburst was roughly 3 days later and leaf elongation was roughly 10 days later).

**Figure 3:**
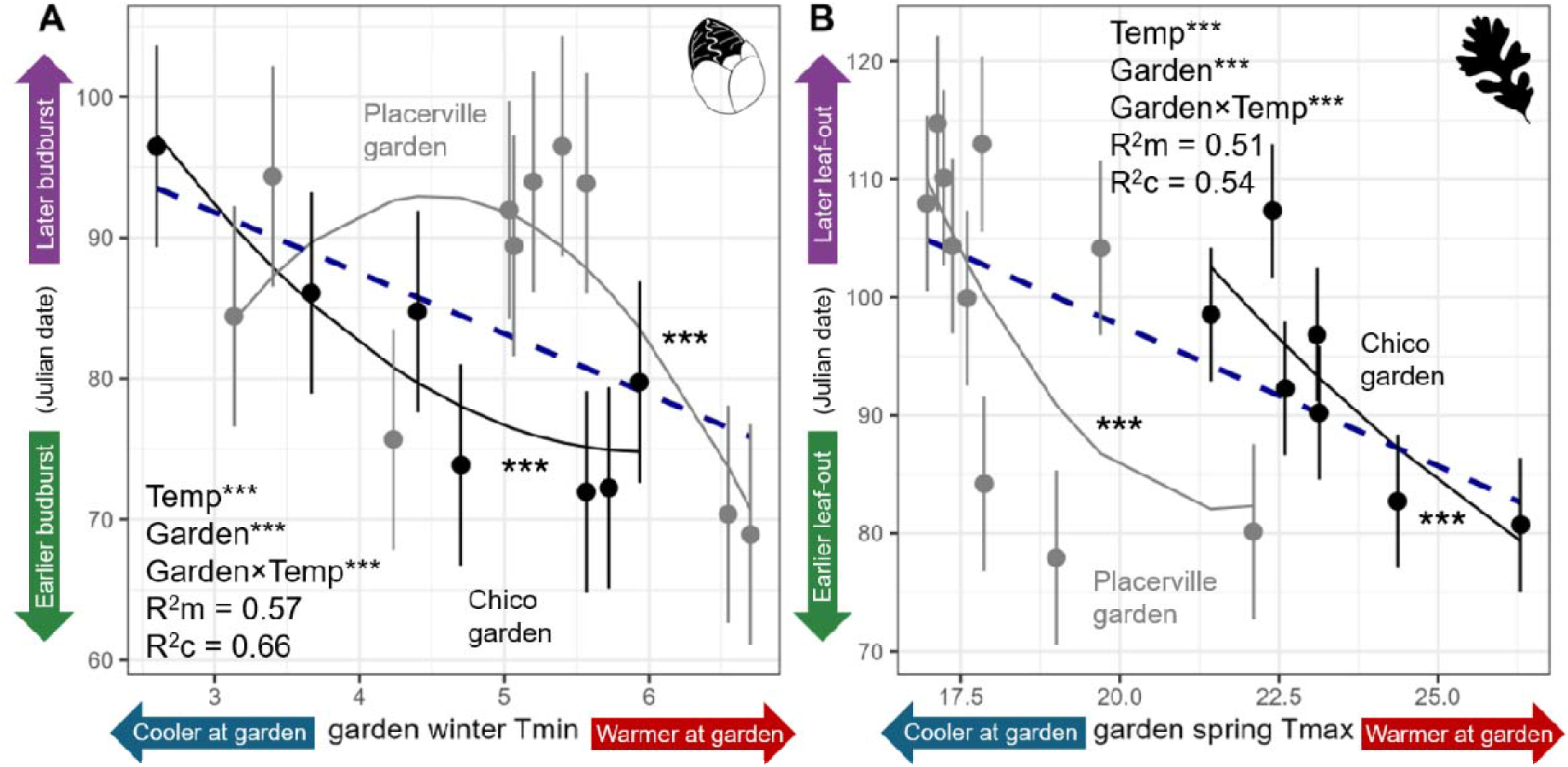
Budburst and leaf elongation show a significant overall trend to emerge earlier in the warmest study years than in the coolest study years. (Blue dashed lines, P < 0.0001, with gardens as replicates). At the Chico garden (black points and lines), the temperature-phenology relationships tended to be more linear than at the Placerville garden (grey points and lines) for both (A) budburst and (B) leaf elongation. Points and error bars show means and standard deviations of best linear unbiased predictors (BLUPs) by year and garden. Black and grey regression lines are from mixed models that incorporated the interaction between garden and temperature. Garden temperature was modeled using a quadratic term in both models. Refer to Tables S7-S8 for model summaries.

### *Quercus lobata* trees track temperature changes

In general, trees burst bud and leaves elongated earlier in warmer years (Figure 3A, 3B, blue dashed lines). Based on a comparison of AIC among models, minimum winter garden temperature (Figure 3A) was the best predictor of budburst date while maximum spring garden temperature (Figure 3B) was the best predictor of leaf elongation date (Table S6). Thus, we used them in all subsequent analyses. The Chico garden (black points and lines) showed a more consistent negative trend between date of bud burst or elongation than the Placerville garden (grey points and lines), indicating that phenological patterns at the Chico garden are much earlier than at the Placerville garden under the same warm temperatures. At the Placerville garden, five of the study years (including the two warmest and coolest years) showed a clear association between temperature and budburst date, but five study years with very similar minimum temperatures (roughly 5.0-5.5°C) had late budburst date (Figure 3A, grey line). These differences in phenological timing indicate that either the much colder winter temperatures at Placerville or the much warmer year-round temperatures at Chico (Figure S1) may influence the date of budburst and leaf elongation to differ at the same spring temperatures.

### Trees from warmer origins respond differently to temperature variation

Despite clear effects of origin temperature on phenology, most trees showed a high capacity to track changes in temperature among years (Figure 4). Warm-origin trees tended to show higher capacity to burst bud earlier in warm years while cool-origin trees showed higher capacity to burst bud later in cool years (Figure 4A). Date of leaf elongation (Figure 4B) showed some evidence of asymmetry in tracking, i.e. the differences between leaf elongation dates of warm-origin trees and cool-origin trees were greater in warmer years than cooler years. This discrepancy suggests that warmer-origin trees are only slightly less capable of delaying phenology in response to cold years relative to cool-origin trees but are much more capable of advancing phenology in response to warm years. As a result, warm-origin trees exhibit greater capacity to lengthen their growing seasons under favorable conditions and only moderately increase their risk of cold damage, while cool-origin trees are more likely to delay phenology longer than necessary.

**Figure 4:**
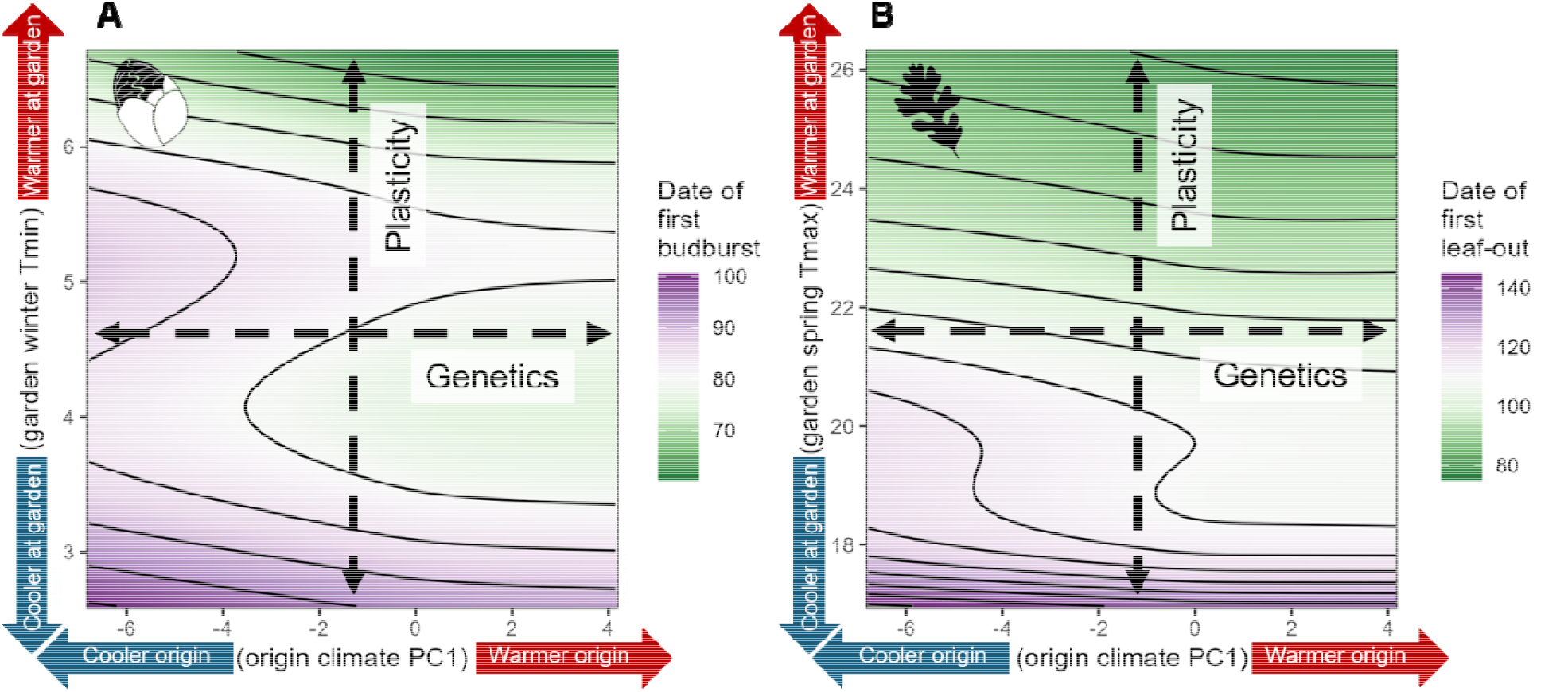
Date of budburst and leaf elongation are affected by an interaction between origin climate and garden temperature fluctuations. Discrepancy in temperature tracking among trees from different climates gives warm-origin trees a slightly longer growing season than cool-origin trees under warm conditions. **A:** Budburst date as a function of climate of origin PC vector (see Methods, “Principal components analysis”) and average monthly minimum garden temperature, Dec-Feb. **B:** Leaf elongation date as a function of climate of origin PC vector and average monthly maximum garden temperature, Mar-May. Refer to Tables S9-S10 for model summaries.

### Phenology is under selection in the garden environments

Date of budburst and leaf elongation showed significant, but directionally variable, effects on relative growth rates in the garden environments (Figure 5). At the Chico garden, we found evidence of stabilizing selection on budburst date, with a significant quadratic relationship between budburst date and relative growth rate (RGR; Figure 5A). Maximum growth was seen in trees with average budburst dates around March 21^st^ (the 80^th^ day of the year). We did not find a significant relationship between budburst date and RGR at the Placerville garden (Figure 5A). By contrast, we observed evidence of directional selection at the Chico garden, and stabilizing selection at the Placerville garden, on date of leaf elongation (Figure 5B). Earlier leaf elongation was associated with higher RGR at the Chico garden, and the highest RGR was observed in trees with an average leaf elongation date around 95 at the Placerville garden. Thus, we see environment-dependent effects of phenology on a component of fitness. Even at the Chico garden, which rarely experiences freezing temperatures during the budburst period, trees showed reduced growth from early budburst.

**Figure 5:**
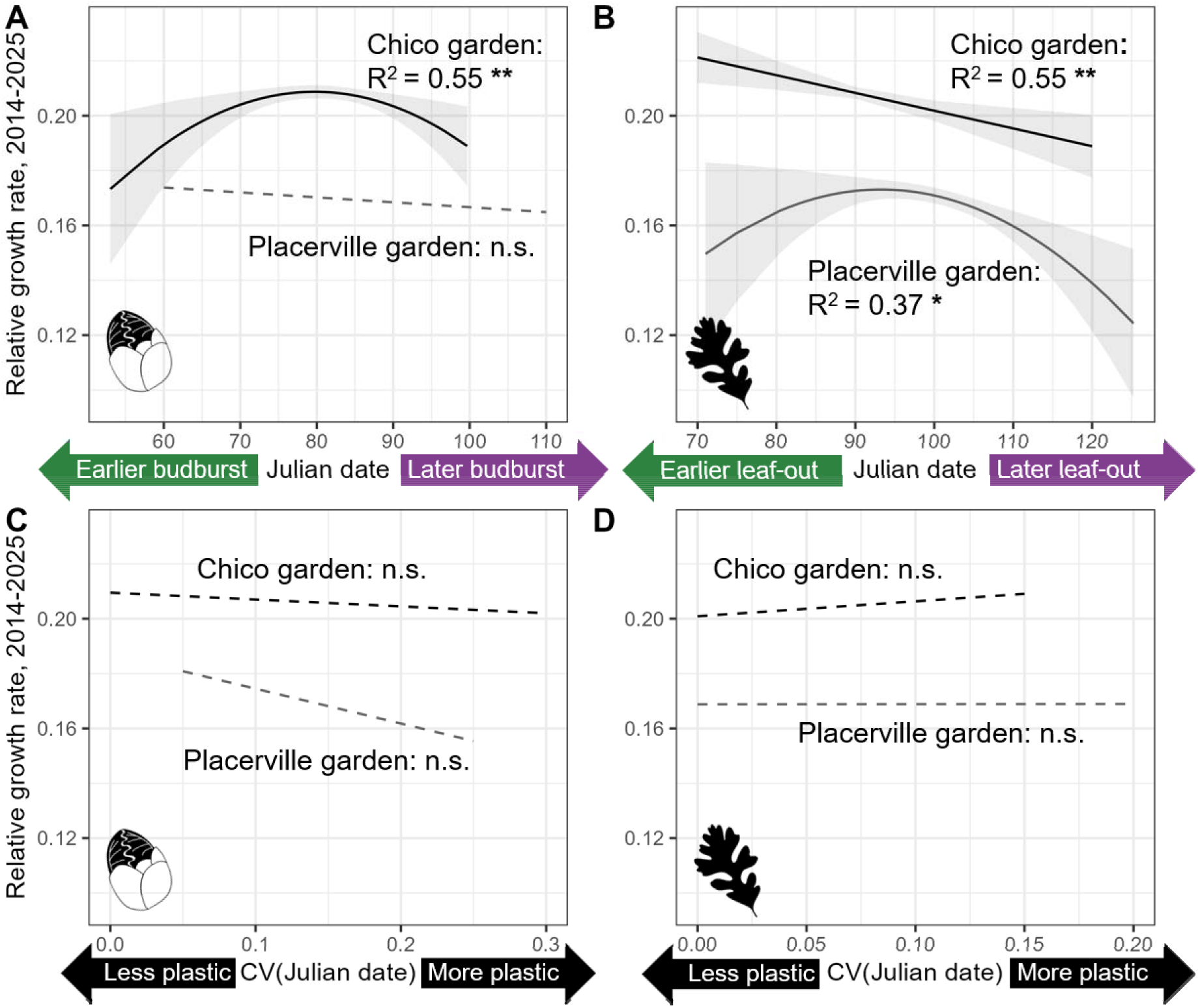
Evidence of environment-dependent stabilizing and directional selection, but no cost to plasticity, on spring leaf phenology in *Quercus lobata*. **A:** At the Chico garden, intermediate dates of first budburst were associated with higher relative growth rate (RGR); no relationship between date of first budburst and cumulative RGR (2014-2025) was observed at the Placerville garden. **B:** At the Chico garden, early leaf elongation dates were associated with higher RGR; at the Placerville garden, intermediate leaf elongation dates were associated with higher RGR. **C:** No relationship between plasticity in budburst date (coefficient of variation among years) and RGR at either garden. **D:** No relationship between degree of plasticity in leaf elongation date (coefficient of variation among years) and RGR at either garden. Dates of budburst and leaf elongation represent averages across years for each tree, thus representing overall relative trends among trees across all study years. Relative growth rates represent individual growth between 2014 and 2025. Linear models accounted for trees’ climate of origin and height prior to outplanting in the gardens. To account for different scales among variables, all independent variables were scaled prior to fitting. Refer to Methods, “Evidence of costs to plasticity” for more detail on the plasticity metric. Refer to Tables S11-S14 for model summaries.

We did not find evidence of a cost or benefit to phenological plasticity among years (Figure 5C, 5D). In addition, we found no effect of the variation in trees’ dates of budburst (Figure 5C) or leaf elongation (Figure 5D) among years on their cumulative RGR when controlling for initial height and climate of origin. However, climate of origin was associated with degree of individual plasticity among years. Trees from hotter, drier climates had greater plasticity in budburst date at the Chico garden (Figure S6A) and greater plasticity in leaf elongation date than trees from cooler, wetter climates at the Placerville garden (Figure S6B).

## DISCUSSION

*Quercus lobata* trees tracked temperature fluctuations through phenological plasticity, which was the primary driver in the timing of budburst and leaf elongation. However, evolutionary history may constrain the extent to which plasticity can maintain optimal phenology under a changing climate. We found evidence that contemporary selection acts on phenology, with directional selection favoring earlier leaf elongation, but not earlier budburst, in the warmer garden environment. However, neither phenological plasticity or earlier budburst was associated with a growth advantage in either garden. Thus, although selection can act on phenological traits, our results provided limited evidence that selection currently favors phenological responses that would enhance growth under warmer conditions. Together, these findings suggest that climate of origin can constrain phenological tracking and that contemporary selection may not be sufficient to overcome these constraints. As temperatures continue to warm, trees adapted to cooler climates may become increasingly mismatched with local environmental conditions.

### Genetic differentiation and climate of origin shapes spring leaf phenology

Spring leaf phenology of *Q. lobata* was genetically differentiated among families sampled from throughout the species range and strongly associated with climate at the site of origin. Trees from warmer, drier environments consistently experienced earlier budburst and leaf elongation than trees from cooler, wetter environments despite interannual temperature fluctuations. This pattern was consistent between the two common gardens, with warm-origin trees bursting bud earlier than cool-origin trees. Therefore, we found strong evidence that historical selection has shaped phenology in relation to climate. This finding that phenological timing is genetically based and associated with climate of seed source has been documented in European oaks (Ducousso et al., 1996; Jensen and Hansen, 2008; Vitasse et al., 2009; Alberto et al., 2011; Ramírez-Valiente et al., 2022) and other forest tree species (Rehfeldt, 1989; Rehfeldt et al., 1999).

The pattern of earlier phenology in warm-adapted trees was observed across 9 study years representing 13 cumulative years of growth, indicating that relationships between environment and phenotype are conserved across the entire juvenile stage of the trees’ development. Our findings extend those of Wright et al. (2021), who found a significant negative relationship between origin temperature and budburst date in the same trees across two seasons (2018 and 2019, when the trees were 6 and 7 years old, respectively). While phenology is often predicted to change as trees age (Vitasse et al., 2013), our results support those of Thomas et al. (2024), who observed differences among subpopulations in budburst dates of 40-year-old *Quercus alba* trees in a common garden in the northeastern United States. The finding that relationships between phenotypes and origin climates in *Q. lobata* establish early and remain consistent throughout the juvenile stage also matches observed growth trends from the same study (Goetz et al., 2026). Therefore, in oaks, the genetic basis of phenology persists throughout the life span with no evidence of complete acclimation to the garden environment.

### Trees from warmer origins have greater capacity to track temperature change

Overall, *Q. lobata* trees showed a high capacity to adjust phenology to temperature fluctuations, but higher plasticity and a tendency toward earlier leaf development gave trees from warmer sites a greater capacity to track interannual temperature fluctuations— i.e., burst bud and elongate leaves earlier during warmer springs. Trees from warmer climates showed greater phenological plasticity, as indicated by the interaction between origin and garden climate on phenology and the association between origin climate and individual coefficients of variation in phenology among years. Differentiation in phenology among localities has been found in several other oak species (Vitasse et al., 2010; Dantec et al., 2014; Dewan et al., 2020; Meger et al., 2024; Wu et al., 2026). In addition, our findings parallel other studies demonstrating that trees from warmer, drier localities tend to show higher plasticity in their traits than trees from cooler, wetter provenances (Nicotra et al., 2010; Lloret et al., 2011; Nicotra et al., 2015), including phenology (Morin et al., 2010; Cooper et al., 2019; Knott et al., 2023), when planted in common gardens. Vitasse et al. (2010), who studied sessile oak and beech populations in Europe, provide the only example where trees responded plastically to warming but local populations did not differ in their degree of plasticity. Here, we generalize these findings by demonstrating empirically that across a range-wide climatic gradient, budburst and leaf elongation dates are predicted by the interaction between a tree’s climate of origin and the temperatures it experiences during the leaf emergence period. Thus, many tree populations that have evolved in cooler climates may have limited capacity to maintain appropriate phenology under climate warming.

Determining the cues that drive species’ phenology is crucial for accurately predicting future shifts (Hänninen et al., 2019; Inouye et al., 2019). Our finding that minimum temperature best explained interannual fluctuations in *Q. lobata* budburst agrees with Gerst et al. (2017), who found the same pattern in naturally occurring populations. Other oak species have also been shown to respond strongly to rising spring temperatures by advancing budburst and leaf elongation (Vitasse et al., 2013; Dantec et al., 2014; Dantec et al., 2015). However, we also found small but significant relationships between latitude of origin and phenology, suggesting an influence of photoperiod (Marchin et al., 2015; Flynn and Wolkovich, 2018). In other species, photoperiod effects can be a major source of constraints to plasticity because photoperiod remains static despite warming temperatures (Way and Montgomery, 2015; Ford et al., 2017). However, within the *Q. lobata* range, latitude and climate are only weakly correlated (Figure S3) because California’s topography supports a wide variety of microclimates within small spatial extents (Sork et al., 2010b; Ackerly et al., 2020). As a result, it is likely that local adaptation to climate drives the observed constraints on plasticity (Alberto et al., 2013). Finally, large differences in phenological patterns between the garden sites demonstrated the influence of the local environment and timing of warming on phenological response, as previously shown in wild populations of *Q. lobata* (Gerst et al., 2017). Trees growing in the Chico garden, which experiences earlier and faster spring warming than the Placerville garden, showed a consistent linear response of earlier phenology in warmer years. By contrast, trees in the Placerville garden had highly variable responses across years, with the latest budburst observed in years with intermediate temperatures. This discrepancy underscores the complex nature of intraspecific variation in phenological cues, especially regarding strategies that minimize temperature-related and biotic risks (Augspurger, 2013; Dantec et al., 2015; Marchin et al., 2015; Silvestro et al., 2019). Therefore, our findings reveal that temperature is a powerful driver of both adaptation and plasticity in *Q. lobata* phenology.

Tracking temperature via phenology may be beneficial to tree performance under climate change, but its benefits may be limited. We found only minor fitness benefits of earlier leaf elongation at the warmer Chico garden, suggesting that adjusting phenology through plasticity or selection is unlikely to mitigate predicted growth declines due to stress caused by high summer temperatures (Goetz et al., 2026). This finding is concordant with other studies demonstrating that tracking can increase the length of the growing season and promote annual growth (Larcher, 2003; Augspurger, 2008), but growth may decline with increased stress later in the season (Knott et al., 2023; Bose et al., 2025). In addition, many plant species can track temperature phenologically only up to a threshold, beyond which phenology becomes mismatched with local climate (Morin et al., 2010; Lapenis et al., 2014; Ford et al., 2017; Lu et al., 2025). Finally, adjusted phenology may increase asynchrony with other organisms, leading to other fitness disadvantages unrelated to temperature stress (e.g., Marquis and Whelan, 1994; Dantec et al., 2015). Therefore, while phenology shapes family responses to climate conditions, adjusted phenology alone will likely not allow trees to maintain their fitness under altered climate conditions.

### Contemporary selection on phenology helps explain maintenance of constraints

Associations between spring phenology and fitness suggested that contemporary selection plays a role in maintaining constraints on phenology. In both gardens, trees exhibited associations between patterns of relative growth rate, a fitness component, and spring phenology, indicating selection on leaf emergence timing. We found evidence of stabilizing selection on date of budburst at the warmer Chico garden—with peak relative growth rates associated with mid-March budburst dates—implying that early budburst is disadvantageous even in the warmer portion of the species range, either due to cold stress or biotic factors (Dantec et al., 2015). Moreover, in the same garden, we found significant directional selection favoring earlier leaf elongation, suggesting a de-coupling of the two developmental states. In contrast, we observed a different pattern of selection at the Placerville garden. There, we found no evidence of selection favoring earlier budburst but did find stabilizing selection on leaf elongation date, with peak relative growth rates in early April. This peak suggests selection against early leaf elongation, possibly because later leaf elongation would reduce the risk of frost damage due to cold temperatures in early spring (Alberto et al., 2011; Dantec et al., 2015). These trends are consistent with, and may explain, our earlier finding that *Q. lobata* shows strong signatures of local adaptation to modern winter temperatures despite maladaptation to summer temperature maxima (Goetz et al., 2026). Thus, we expect that, under range-wide warming, trees adapted to warmer climates may have a growth advantage because they have greater capacity to advance their phenology than trees adapted to cooler climates. However, selection against early phenology in both environments demonstrates a means by which constraints to phenology are maintained, even as temperatures rise.

Plasticity can weaken selection if it allows trees to respond beneficially to temperature fluctuations (Sultan, 1987), but its evolution may be constrained when the cost of adjusting physiological processes reduces a fitness component, such as growth (DeWitt et al., 1998). We found no evidence that plasticity conferred a growth benefit, nor evidence of a cost, indicating that our previous finding that warmer-origin trees grew more than cooler-origin trees in the common gardens (Goetz et al., 2026) is likely explained by traits other than phenological tracking capacity. Thus, our results provide no evidence that selection has favored increased plasticity in *Q. lobata* beyond its existing response to warming (Crispo et al., 2010). Positive effects of plasticity, therefore, may not increase over time, as predicted or observed in other tree species (Ghalambor et al., 2007; Cleland et al., 2012; Cooper et al., 2019; Wang et al., 2023). By contrast, selection on the genetic mechanisms underlying phenology may shift allele frequencies over time (Körner and Basler, 2010; Zohner and Renner, 2014). For example, sessile oak, *Quercus petraea*, shows a high capacity to rapidly evolve phenological changes in response to warming (Alberto et al., 2013; Caignard et al., 2024; Losch et al., 2025), but other oak species appear to be more constrained in their evolutionary responses to warming (Knott et al., 2023; Thomas et al., 2024). Given evidence that *Q. lobata* trees exhibit an adaptational lag to warming (Browne et al., 2019; Goetz et al., 2026), it is unknown whether selection can increase temperature-tracking capacity.

### Conclusions

The degree to which plants can respond to environmental fluctuations is determined in part by natural selection on inherited traits and by the capacity of plants to adjust these traits in response to altered conditions. Our findings suggest that *Q. lobata* populations adapted to warmer climates will likely be able to adjust their phenology to keep up with rising temperatures, which may help maintain phenological synchrony with increasingly warm spring conditions. By contrast, phenology in populations adapted to cooler climates will likely become increasingly decoupled from timing of spring temperature increases. However, although *Q. lobata* populations can track interannual temperature variation through phenology, plasticity does not promote higher growth and thus may not be sufficient to buffer populations from continued warming. Given that cooler-origin trees are also more vulnerable to climate fluctuations than warmer-origin trees (Goetz et al., 2026), their reduced capacity for phenological adjustment may increase climate maladaptation. Cooler-origin populations may therefore experience increasingly mismatched phenology as spring temperatures advance beyond the range over which their plastic responses can compensate. If plasticity does not translate into greater growth, these populations may be less able to buffer the demographic consequences of continued warming, potentially increasing fitness differences among local populations across the species’ range.

## Author contributions

VLS and JWW designed the common garden phenology study and secured funding. All authors contributed to data collection and project management. VLS, JWW, and ARBG designed research questions. ARBG analyzed data and created visualizations with input from VLS. ARBG and VLS wrote the initial draft of the paper, and all authors contributed revisions.

## Data availability

Upon publication, all data and R scripts underlying analyses in this paper will be made available on FigShare (https://doi.org/10.6084/m9.figshare.33302238). In addition, raw data will be made public through the project’s interactive data portal (sorklab2.eeb.ucla.edu/app/).

## Acknowledgements

We acknowledge the native peoples of California as the traditional caretakers of the ecosystems from which these data were collected. We thank Annette Delfino Mix, Lisa Crane, Robin Scibillio, and Marie McLaughlin for their assistance in establishing and maintaining the valley oak common garden experiment. We thank D. Alvarado, M. Antonelli, O. Arellano, R. Belmonte, G. Birdwell, R. Boynton, C. Brackett, N. Burdeinii, C. Cantrell, W. Carlson, N. Corley, A. Curry-Long, S. David, M. Dougherty, E. Estrada, H. French, J. Fucigna, J. Garcia, S. Gribble, A. Guzman, H. Hardie, N. Headley, A. Hickox, A. Kelley, A. King, S. Krasnobrod, A. Lawrence, A. Lecitona, D. Linville, D. Lomeli, A. Moussali, P. Munson, L. Poland, C. Raether, H. Ramirez, A. Reyes, R. Robinson, R. Schafer, L. Schubert, K. Shapiro, FNU Shatakashi, K. Spratt, A. Sullivan, J. Syndor, M. Taylor, S. Taylor, J. Vang, A. Vasanthan, J. Waian, M. Wasim, K. Weycker, and E. Wilkinson for field assistance. We thank Kara Sanghera, Randy Meyer, Mitchell Washington, Thomas Harbin, and Rebecca Cutforth for their assistance with supervising data collection. Finally, we thank L. Peck, R. Buck, B. Badillo, H. Yang, and J. Holmes for substantive feedback on the manuscript. This research was supported for ten years by funds from the USDA Forest Service Pacific Southwest Research Station, CALFIRE agreement 8CA04059, and UCLA. Starting in 2023, ARBG and the project have been supported by the National Science Foundation (NSF-LTREB-2232794), and the Forest Service has provided continued staff support. The findings and conclusions in this publication are those of the authors and should not be construed to represent any official USDA or U.S. Government determination or policy. Any use of product names is for informational purposes only and does not imply endorsement by the U. S. Government.

