## Supplementary information for "Climate of origin constrains phenological tracking of temperature in a California endemic oak (*Quercus lobata*)"

### Supplementary figures

**Figure S1**: Time series of mean daily maximum and minimum temperatures (colored lines) and budburst date (grey rectangles) at the *Quercus lobata* common gardens at the Chico Seed Orchard, Chico, CA (“Chico garden”) and the Institute of Forest Genetics, Placerville, CA (“Placerville garden”). Temperatures were measured from February through May each year and averaged over the duration of the study period (2014-2026). Black vertical lines represent median dates of budburst (first bud emergence) and leaf elongation (full leaf emergence) of *Q. lobata* trees planted at the gardens. Grey rectangles represent the ranges of budburst and leaf elongation dates. Temperature data were acquired from PRISM (prism.oregonstate.edu) by extracting point data from 800m-resolution rasters with interpolation between grid cells.

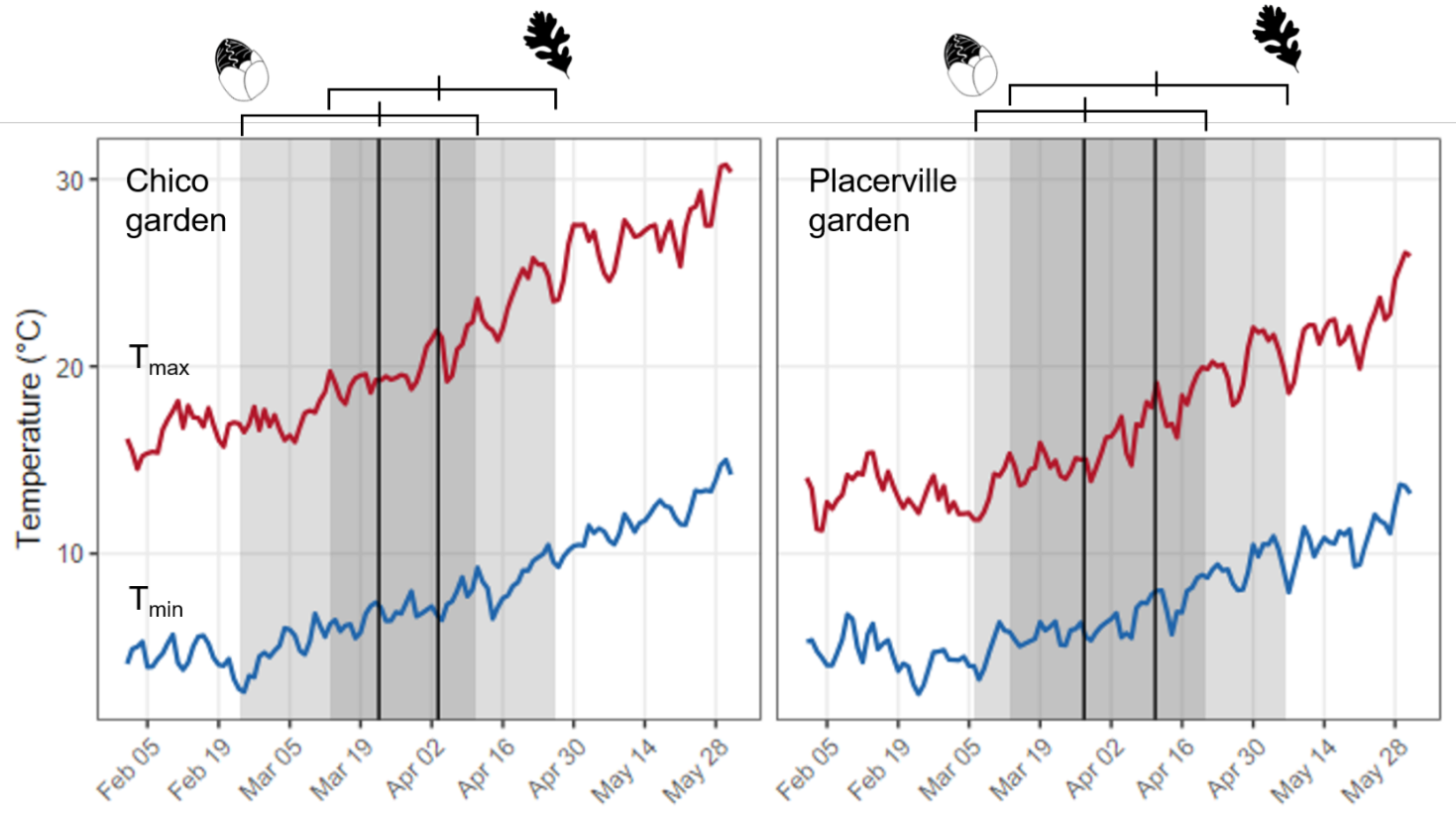

**Figure S2**: **PRISM temperature data accurately reflects conditions in the Chico and Placerville garden sites.** Black lines represent monthly averages of minimum temperatures acquired from the PRISM climate database (prism.oregonstate.edu) for garden point locations, at 800m resolution with grid interpolation, 2014-2025. Red lines represent monthly averages of minimum temperatures measured in the garden plots using iButton temperature sensors, 2015-2019. PRISM data is very strongly correlated with iButton data (R^2^ = 0.98, β = 1.01). Most deviations between data sources occurred at the Chico garden during the summer months, which were not part of the study period. During the study period (shown in blue shading), all deviations were within 1.5°C.

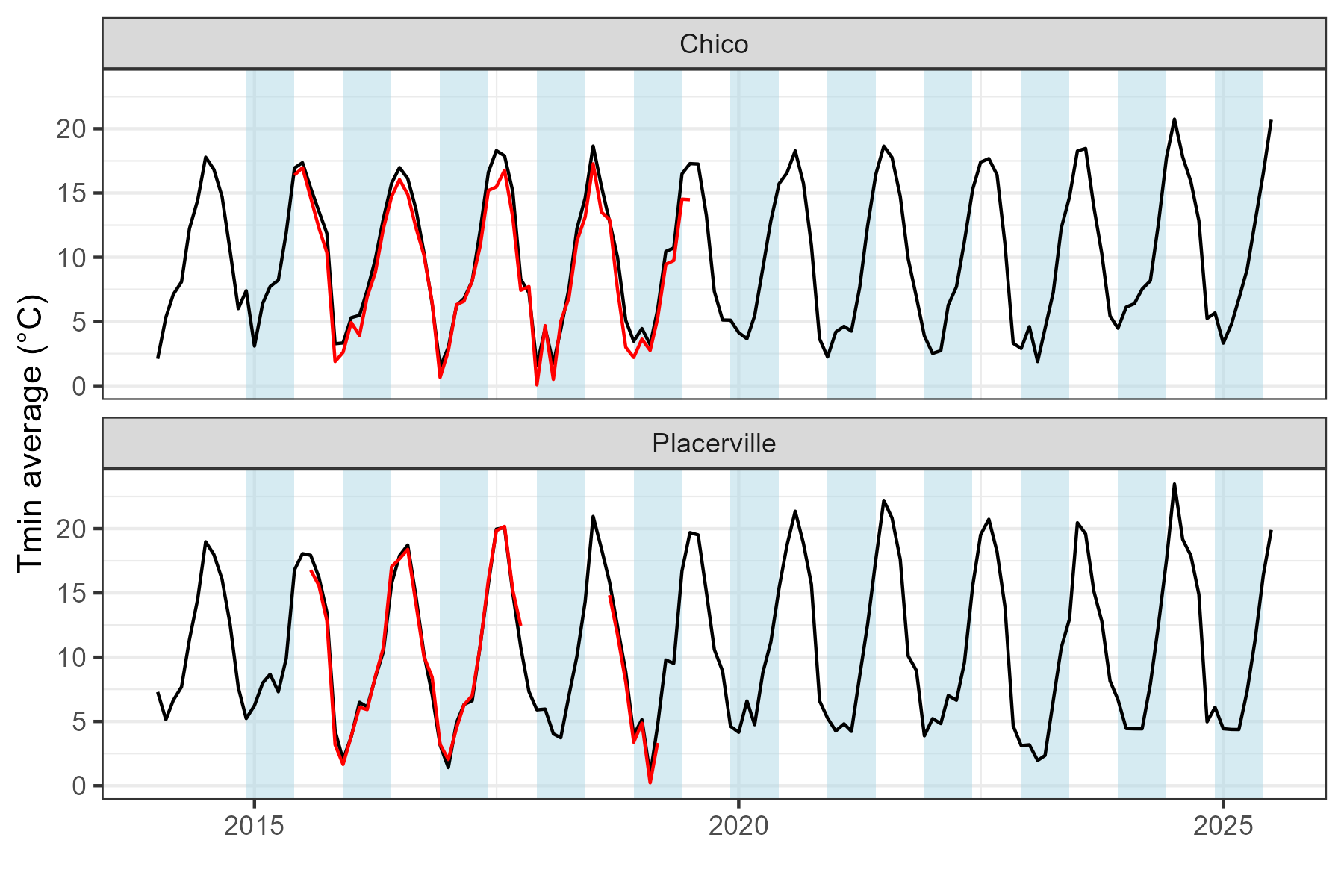

**Figure S3a**: **Significant linear relationship between latitude and climate at point locations of 659 adult Q*uercus lobata* trees whose progeny were planted into two common gardens.** Hotter, drier sites (higher values of PC1 vector) are found at more southern latitudes, and cooler, wetter sites (lower values of PC1 vector) are found at more northern latitudes. PC1 explained 41.1% of variance in climate variables; loadings are shown in **Figure S3b.**

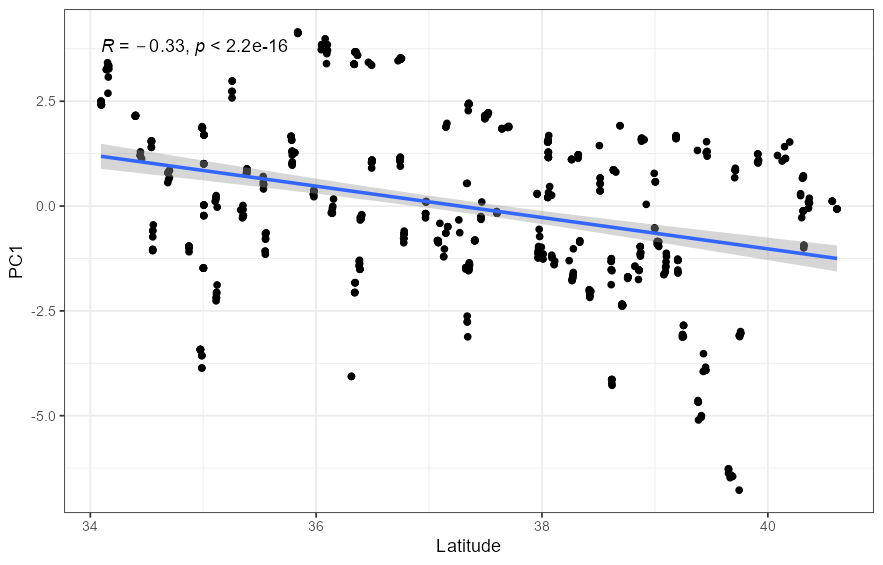

**Figure S3b**: **Biplot showing loadings of the climate variables for the first two PC vectors** (adapted from Browne *et al.* 2019). The climate variables used to characterize climate of origin were maximum summer temperature (Tmax), average maximum temperature across all months (Tmax_annual), minimum winter temperature (Tmin), average minimum temperature across all months (Tmin_annual), average temperature across all months (Tave), temperature seasonality (Bioclim 4), precipitation seasonality (Bioclim 15), summer precipitation (Bioclim 18), precipitation of the coldest quarter (Bioclim 19), and climatic water deficit (CWD). Refer to Methods, “Principal components analysis of origin climate” for more details. Variable loadings are reported in **Table S2**.

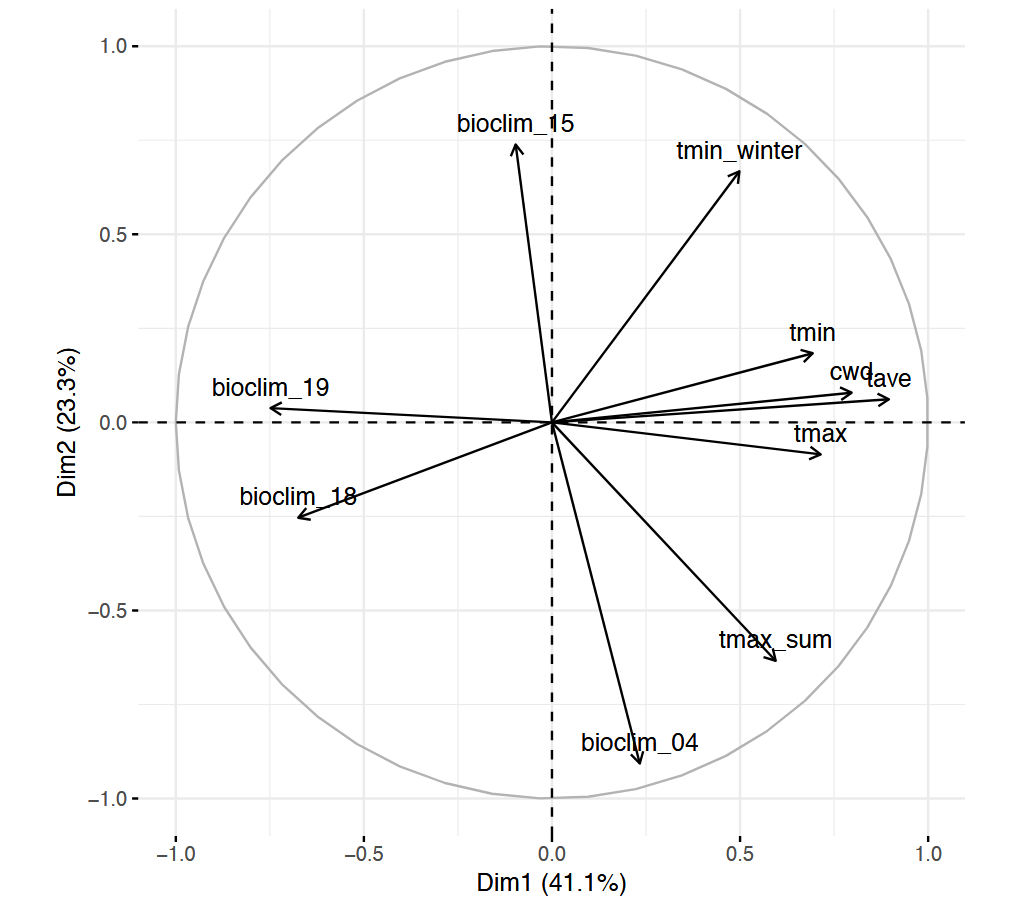

**Figure S4**: B**udburst** **and** **leaf elongation dates** differ among *Q. lobata* families (genotype effect) and between common gardens (environment effect), but do not exhibit an interaction between family and garden (genotype × environment effect). All trees tend to have earlier budburst and leaf elongation date at the Chico garden than at the Placerville garden. Lines are colored by origin climate: Trees from warmer climates (red lines) tend to have earlier budburst and leaf elongation date than trees from cooler climates (blue lines). Refer to **Table S3** for model summary.

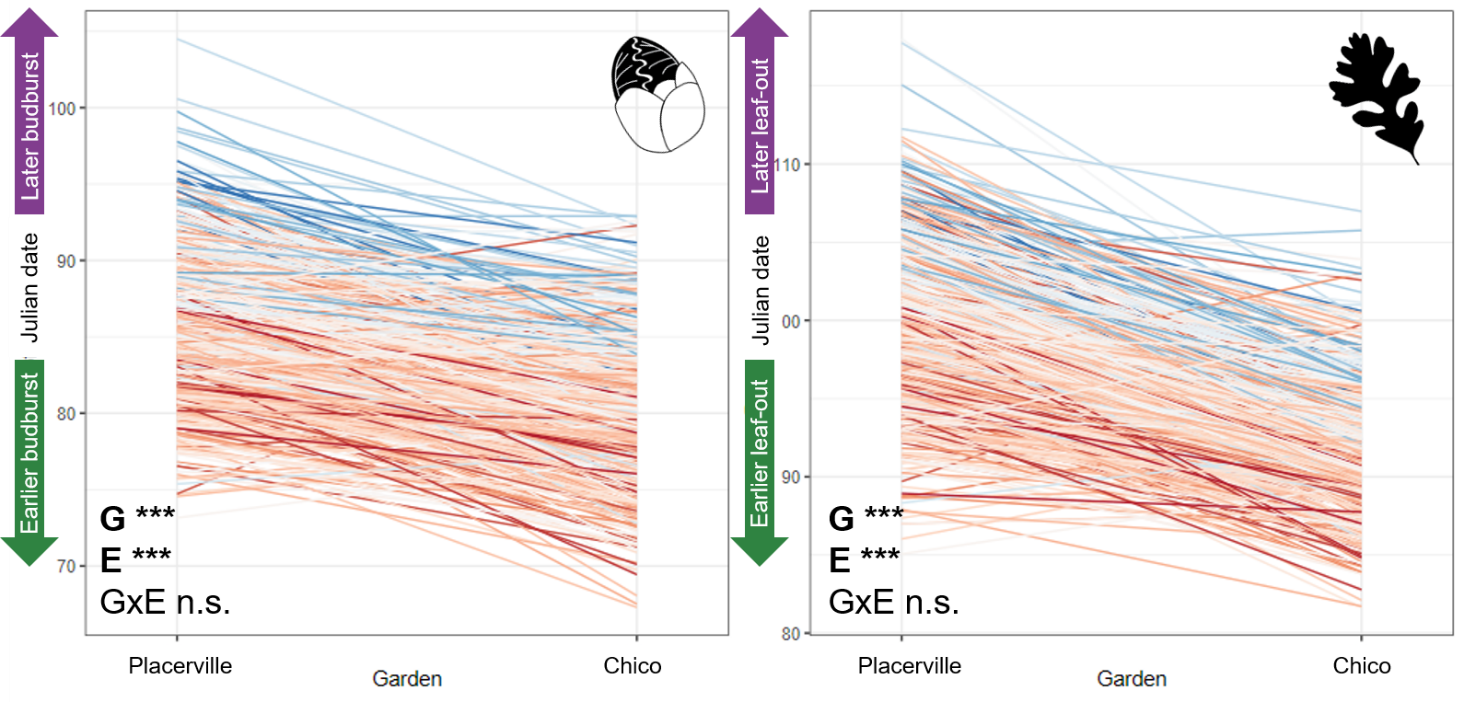

**Figure S5**: **Latitude of origin affects’ (A) budburst and (B)** **leaf elongation dates of *Q.*** ***lobata* trees in two common gardens**. Trees from the northernmost origins burst bud and leafed out roughly 2.5 days earlier than trees from the southernmost origins, suggesting a small effect of origin photoperiod on budburst and leaf elongation date. Models also accounted for garden, origin climate, study year, and the garden*climate interaction.

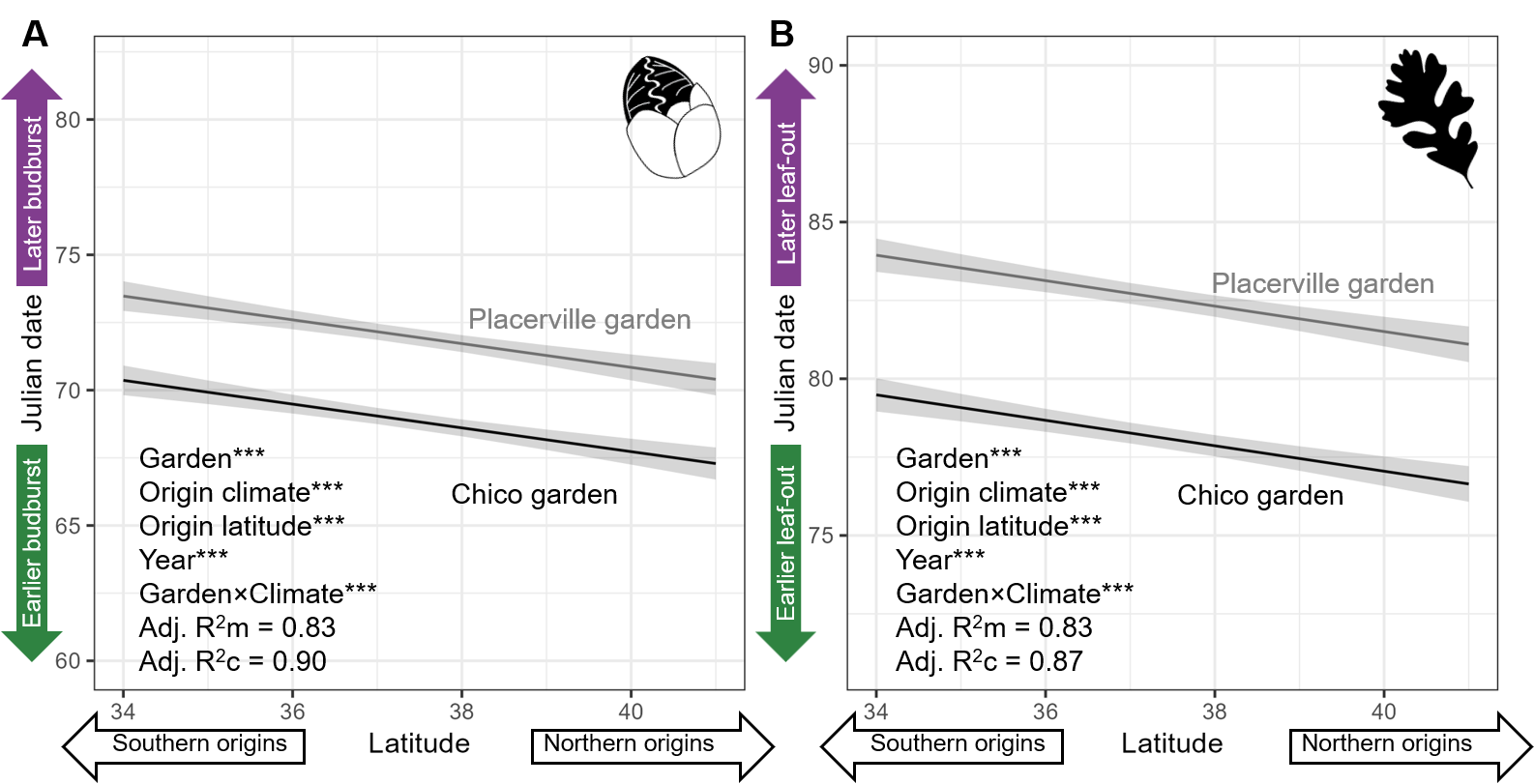

**Figure S6: Climate of origin affects plasticity in (A) budburst and (B) leaf elongation dates of *Q. lobata* trees in two common gardens.** Plasticity was measured as the coefficient of variation of each tree’s average budburst or leaf elongation date over the study period. Trees from hotter, drier climates had greater plasticity in budburst date at the Chico garden (*F* = 7.91, *β* = 0.004, df = 624, *p* = 0.005) and greater plasticity in leaf elongation date than trees from cooler, wetter climates at the Placerville garden (*F* = 8.85, *β* = 0.003, df = 621, *p* = 0.003).

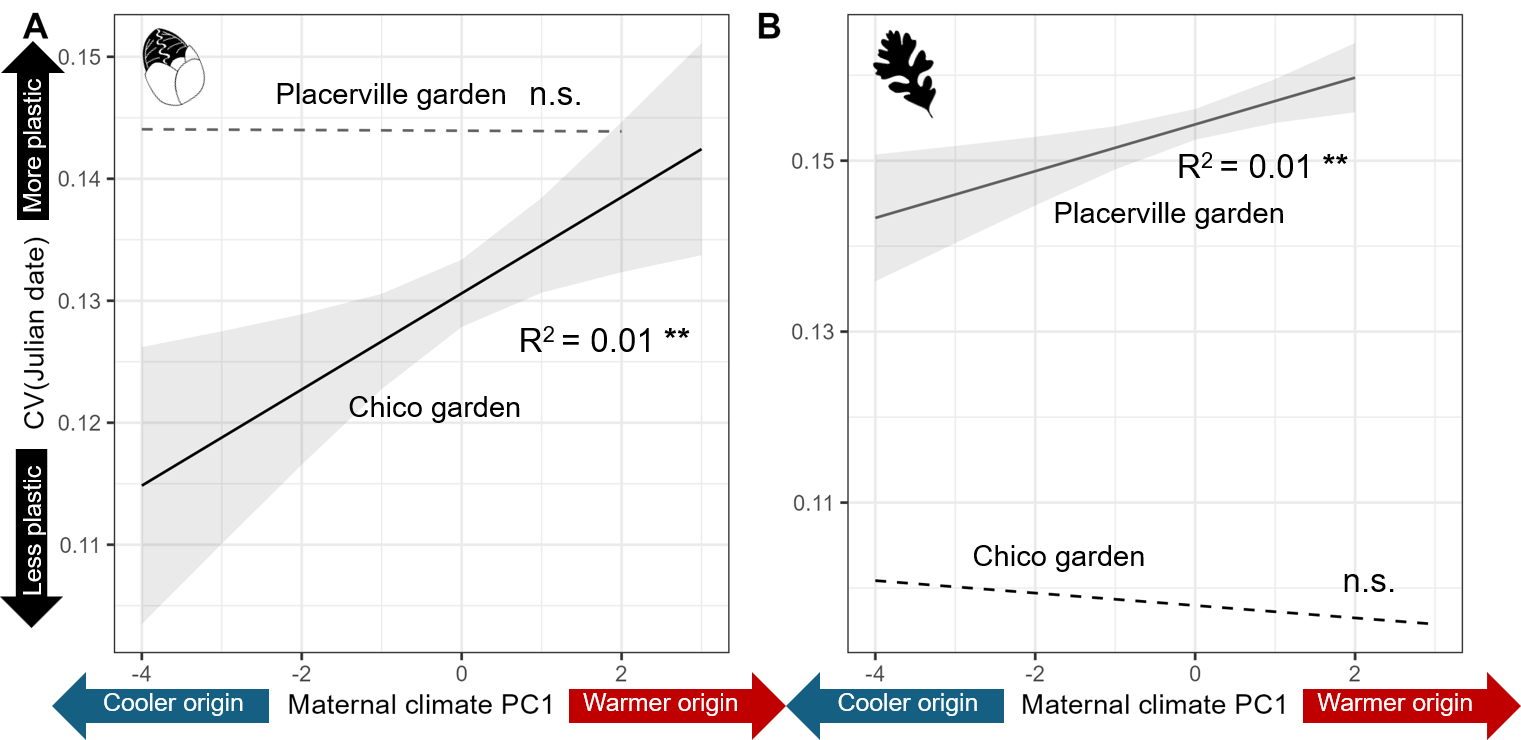

### Supplementary tables

**Table S1**: Observers, survey date ranges, and sample sizes for measurements of **budburst** and **leaf elongation** **dates of *Q. lobata* trees** in two common gardens, 2015-2026. CC: C. Canning; AG: A. Goetz. The “Multiple” observer label refers to two cohorts of independent student researchers whose phenological measurements were found to significantly differ (Wright *et al.* 2021), so they are counted separately in mixed models. The “AG” label also refers to cohorts of student researchers, but they were closely supervised for consistent data collection throughout the study period and were haphazardly assigned to survey multiple areas of the plots to remove any observer effect.

| Garden | Year | Observer | Survey date range | N_Budburst_ | N_Leaf elongation_ |
| --- | --- | --- | --- | --- | --- |
| Placerville | 2015 | CC | Feb 4-Mar 31 | 1632 | 1600 |
|  | 2016 | CC | Mar 8-Apr 14 | 1628 | 1321 |
|  | 2017 | CC | Mar 20-Apr 26 | 1610 | 899 |
|  | 2018 | CC | Feb 12-May 11 | 1651 | 1649 |
|  | 2019 | CC | Mar 20-May 11 | 1652 | 1649 |
|  | 2020 | CC | Mar 10-May 13 | 1128 | 548 |
|  | 2021 | CC | Mar 26-May 6 | 799 | 784 |
|  | 2024 | AG | Feb 27-Apr 26 | 1612 | 1495 |
|  | 2025 | AG | Feb 17-Apr 25 | 1626 | 1596 |
|  | 2026 | AG | Feb 27-Apr 10 | 1129 | 868 |
| Chico | 2015 | CC | Feb 2-Apr 2 | 1633 | 1283 |
|  | 2016 | CC | Feb 23-Apr 7 | 848 | 670 |
|  | 2018 | Multiple | Feb 11-May 2 | 1098 | 863 |
|  | 2019 | Multiple | Feb 9-Apr 30 | 1684 | 1598 |
|  | 2024 | AG | Feb 27-Apr 19 | 1682 | 1556 |
|  | 2025 | AG | Feb 11-Apr 22 | 1696 | 1664 |
|  | 2026 | AG | Feb 24-Apr 3 | 1982 | 1941 |

**Table S2**: Loadings of variables on first principal component vector of variation in climate among 673 *Q. lobata* trees used as seed sources in the common garden study.

| Variable | PC1 loading |
| --- | --- |
| Tmax_sum: Summer maximum temperature (Jun/Jul/Aug avg.) | 0.32 |
| Tmax: Annual maximum temperature (12-month avg.) | 0.38 |
| Tmin: Annual minimum temperature (12-month avg.) | 0.31 |
| Tave: (Tmax + Tmin) / 2 | 0.44 |
| Tmin_winter: Winter average minimum temperature (Dec/Jan/Feb avg.) | 0.22 |
| CWD: Climatic water deficit | 0.39 |
| bioclim_04: Temperature seasonality | 0.12 |
| bioclim_15: Precipitation seasonality | -0.04 |
| bioclim_18: Precipitation of warmest quarter | -0.34 |
| bioclim_19: Precipitation of coldest quarter | -0.37 |

**Table S3**: **Type III repeated-measures ANCOVA tests comparing interannual variation in budburst (A) and leaf elongation (B) date** among families of *Q. lobata* trees growing in two common gardens. Budburst and leaf elongation date significantly differed among families and years, with a significant interaction between family and year.

|  | | A. Budburst date | | | | B. Leaf elongation date | | | |
| --- | --- | --- | --- | --- | --- | --- | --- | --- | --- |
|  | *df_num_* | | *df_den_* | *F* | *p* | *df_num_* | *df_den_* | *F* | *p* |
| Year | | 8 | 21,352.61 | 2,546.14 | **< 0.001** | 8 | 18,566.69 | 4,441.35 | **< 0.001** |
| Garden | | 1 | 21,588.31 | 782.00 | **< 0.001** | 1 | 18,759.64 | 2,181.89 | **< 0.001** |
| Year × Garden | | 5 | 21,346.40 | 471.04 | **< 0.001** | 5 | 18,552.83 | 1,513.43 | **< 0.001** |

**Table S4**: **Results of repeated-measures ANCOVA tests comparing average** **(A) budburst and (B) leaf elongation date** among families of *Q. lobata* trees growing in two common gardens. Model also accounted for the fixed effect of study year. Both budburst and leaf elongation date significantly differed between gardens and among families, but the Garden × Family interaction term was not significant.

|  | A: Budburst date | | | B: Leaf elongation date | | |
| --- | --- | --- | --- | --- | --- | --- |
|  | *df* | *F* | *p* | *df* | *F* | *p* |
| Garden | 1 | 972.15 | **< 0.001** | 1 | 2,093.64 | **< 0.001** |
| Family | 620 | 4.08 | **< 0.001** | 620 | 3.71 | **< 0.001** |
| Garden × Family | 620 | 1.02 | 0.37 | 620 | 1.02 | 0.35 |
| Residuals | 2,098 |  |  | 2,096 |  |  |

**Table S5**: Summary of repeated-measures linear mixed model testing the effect of **origin climate (PC1) and garden on (A) budburst and (B) leaf elongation date**. The model also accounted for latitude of origin and study year. The random effect of family was included to account for the fact that these were repeated measurements on the same trees. Dates of budburst and leaf elongation were estimated using BLUPs (see Methods, “Best linear unbiased predictors”). σ^2^: Random effect within-group variance; τ_00_: Random effect between-group variance, ICC: Intra-class correlation coefficient. Random effect variances and *R*^2^ values are estimated using the method of Nakagawa *et al.* (2017).

|  | A. **Budburst date** | | | | | B. **Leaf elongation date** | | |
| --- | --- | --- | --- | --- | --- | --- | --- | --- |
| *Predictors* | *Estimates* | | *CI* | | *p* | *Estimates* | *CI* | *p* |
| (Intercept) | 85.26 | | 80.06 – 90.46 | | **<0.001** | 93.27 | 88.56 – 97.97 | **<0.001** |
| Garden [Placerville] | 3.11 | | 2.97 – 3.25 | | **<0.001** | 4.45 | 4.26 – 4.64 | **<0.001** |
| Climate PC1 | -0.83 | | -0.95 – -0.70 | | **<0.001** | -0.63 | -0.76 – -0.51 | **<0.001** |
| Latitude | -0.44 | | -0.58 – -0.30 | | **<0.001** | -0.41 | -0.53 – -0.28 | **<0.001** |
| Year [2016] | 4.31 | | 4.06 – 4.57 | | **<0.001** | 6.80 | 6.45 – 7.15 | **<0.001** |
| Year [2017] | 12.41 | | 12.09 – 12.72 | | **<0.001** | 17.34 | 16.89 – 17.78 | **<0.001** |
| Year [2018] | 24.73 | | 24.47 – 24.98 | | **<0.001** | 27.28 | 26.93 – 27.63 | **<0.001** |
| Year [2019] | 19.75 | | 19.50 – 20.00 | | **<0.001** | 26.23 | 25.88 – 26.58 | **<0.001** |
| Year [2020] | 24.52 | | 24.20 – 24.84 | | **<0.001** | 30.40 | 29.96 – 30.85 | **<0.001** |
| Year [2021] | 19.91 | | 19.58 – 20.25 | | **<0.001** | 21.48 | 21.02 – 21.95 | **<0.001** |
| Year [2024] | 16.37 | | 16.11 – 16.62 | | **<0.001** | 20.79 | 20.44 – 21.14 | **<0.001** |
| Year [2025] | 16.63 | | 16.38 – 16.88 | | **<0.001** | 20.21 | 19.86 – 20.56 | **<0.001** |
| Year [2026] | 0.97 | | 0.71 – 1.22 | | **<0.001** | 0.19 | -0.16 – 0.54 | 0.293 |
| Garden [Placerville] × PC1 | -0.08 | | -0.15 – -0.02 | | **0.009** | -0.11 | -0.19 – -0.02 | **0.015** |
| **Random Effects** | | | | | | |  |  |
| σ^2^ | | 10.48 | | 20.04 | | |  |  |
| τ_00_ | | 8.36 _Family_ | | 6.13 _Family_ | | |  |  |
| ICC | | 0.44 | | 0.23 | | |  |  |
| N | | 632 _Family_ | | 632 _Family_ | | |  |  |
| Observations | | 10490 | | 10490 | | |  |  |
| Marginal R^2^ / Conditional R^2^ | | 0.826 / 0.903 | | 0.830 / 0.870 | | |  |  |

**Table S6**: **Minimum temperature** (Tmin) best explained variation in **budburst date** and **maximum temperature** (Tmax) best explained variation in **leaf elongation date** in *Q. lobata* trees growing in two common gardens. Linear models were fitted using the family-level BLUPs of budburst and leaf elongation date as the dependent variable and the environmental variable and garden site as independent variables. Models were compared using Akaike’s Information Criterion (AIC) to determine which environmental variable best explained the variation in budburst and leaf elongation date. The random effect of family was included in the model to account for the fact that the same families were compared across multiple years. In subsequent analyses, we tested the effects of minimum temperature on budburst date and maximum temperature on leaf elongation date.

| Phenology variable | Environmental variable | AIC |
| --- | --- | --- |
| Budburst | Precipitation | 77,480.50 |
|  | Tmin | 74,240.60 |
|  | Tmax | 77,599.77 |
| Leaf elongation | Precipitation | 76,851.45 |
|  | Tmin | 77,412.15 |
|  | Tmax | 75,740.62 |

**Table S7**: Summary of linear mixed model testing the effects of **minimum garden temperature on (A)** **budburst date** and **maximum garden temperature on (B) leaf elongation date**. Each analysis combined data from both gardens. Garden temperature was modeled using a linear term. σ^2^: Random effect within-group variance; τ_00_: Random effect between-group variance, ICC: Intra-class correlation coefficient. Random effect variances and *R*^2^ values are estimated using the method of Nakagawa *et al.* (2017).

|  | A. **Budburst date** | | | B. **Leaf elongation date** | | |
| --- | --- | --- | --- | --- | --- | --- |
| *Predictors* | *Estimates* | *CI* | *p* | *Estimates* | *CI* | *p* |
| (Intercept) | 104.72 | 103.96 – 105.49 | **<0.001** | 145.33 | 143.95 – 146.70 | **<0.001** |
| T_min_ winter | -4.31 | -4.45 – -4.16 | **<0.001** |  |  |  |
| T_max_ spring |  |  |  | -2.38 | -2.45 – -2.32 | **<0.001** |
| **Random Effects** | | | | | | |
| σ^2^ | 76.82 | | | 102.88 | | |
| τ_00_ | 7.33 _Family_ | | | 3.21 _Family_ | | |
| ICC | 0.09 | | | 0.03 | | |
| N | 632 _Family_ | | | 632 _Family_ | | |
| Observations | 10490 | | | 10490 | | |
| Marginal R^2^ / Conditional R^2^ | 0.222 / 0.290 | | | 0.313 / 0.334 | | |

**Table S8a**: Summary of linear mixed model testing the **effects of minimum garden temperature on budburst date**, separately by garden. Garden temperature was modeled using a quadratic term. The interaction between garden and temperature was included to test whether interannual temperature differences affected phenology differently at each garden site. σ^2^: Random effect within-group variance; τ_00_: Random effect between-group variance, ICC: Intra-class correlation coefficient. Random effect variances and *R*^2^ values are estimated using the method of Nakagawa *et al.* (2017).

|  | **Budburst date** | | |
| --- | --- | --- | --- |
| *Predictors* | *Estimates* | *CI* | *p* |
| (Intercept) | 79.84 | 79.53 – 80.14 | **<0.001** |
| Tmin winter [1st degree] | -613.49 | -636.03 – -590.94 | **<0.001** |
| Tmin winter [2nd degree] | 282.32 | 259.25 – 305.39 | **<0.001** |
| Garden [Placerville] | 6.53 | 6.29 – 6.77 | **<0.001** |
| Tmin winter [1st degree] × Garden [Placerville] | 446.22 | 417.82 – 474.61 | **<0.001** |
| Tmin winter [2nd degree] × Garden [Placerville] | -949.05 | -977.47 – -920.64 | **<0.001** |
| **Random Effects** | | | |
| σ^2^ | 36.75 | | |
| τ_00_ _Family_ | 9.58 | | |
| ICC | 0.21 | | |
| N _Family_ | 632 | | |
| Observations | 10490 | | |
| Marginal R^2^ / Conditional R^2^ | 0.571 / 0.660 | | |

**Table S8b**: **Summary of linear mixed model testing the** **effects of maximum garden temperature on leaf elongation date**, separately by garden. Garden temperature was modeled using a quadratic term. The interaction between garden and temperature was included to test whether interannual temperature differences affected phenology differently at each garden site. σ^2^: Random effect within-group variance; τ_00_: Random effect between-group variance, ICC: Intra-class correlation coefficient. Random effect variances and R^2^ values are estimated using the method of Nakagawa *et al.* (2017).

|  | **Leaf elongation date** | | |
| --- | --- | --- | --- |
| *Predictors* | *Estimates* | *CI* | *p* |
| (Intercept) | 111.35 | 109.17 – 113.53 | **<0.001** |
| Tmax spring [1st degree] | -1863.42 | -2072.81 – -1654.03 | **<0.001** |
| Tmax spring [2nd degree] | 172.41 | 92.36 – 252.46 | **<0.001** |
| Garden [Placerville] | -16.20 | -18.46 – -13.93 | **<0.001** |
| Tmax spring [1st degree] × Garden [Placerville] | 1305.54 | 1075.47 – 1535.61 | **<0.001** |
| Tmax spring [2nd degree] × Garden [Placerville] | 776.96 | 670.43 – 883.49 | **<0.001** |
| **Random Effects** | | | |
| σ^2^ | 70.90 | | |
| τ_00_ _Family_ | 5.17 | | |
| ICC | 0.07 | | |
| N _Family_ | 632 | | |
| Observations | 10490 | | |
| Marginal R^2^ / Conditional R^2^ | 0.508 / 0.541 | | |

**Table S9**: Summary of generalized additive model testing the **effect of the interaction between origin climate and garden temperature on budburst date** in *Q. lobata*. Model also accounted for the effect of garden site. The interaction term was modeled using a tensor spline (Wood 2017).

|  | **Budburst date** | | |
| --- | --- | --- | --- |
| *Predictors* | *Estimates* | *CI* | *p* |
| (Intercept) | 78.58 | 78.32 – 78.84 | **<0.001** |
| Garden [Placerville] | 1.11 | 1.10 – 1.11 | **<0.001** |
| interaction(Tmin winter,PC1) |  |  | **<0.001** |
| Smooth term (Family, random effect) |  |  | **<0.001** |
| Observations | 10490 | | |
| R^2^ | 0.648 | | |

**Table S10:** Summary of generalized additive model testing the **effect of the interaction between origin climate and garden temperature on leaf elongation date** in *Q. lobata*. Model also accounted for the effect of garden site. The interaction term was modeled using a tensor spline (Wood 2017).

|  | **Leaf elongation date** | | |
| --- | --- | --- | --- |
| *Predictors* | *Estimates* | *CI* | *p* |
| (Intercept) | 107.48 | 106.95 – 108.01 | **<0.001** |
| Garden [Placerville] | 0.83 | 0.82 – 0.84 | **<0.001** |
| interaction(Tmax spring,PC1) |  |  | **<0.001** |
| Smooth term (Family) |  |  | **<0.001** |
| Observations | 10490 | | |
| R^2^ | 0.584 | | |

**Table S11**: Summary of linear mixed model testing the **effects of budburst date on cumulative 2014-2025 relative growth rates** in *Q. lobata t*rees growing in two common gardens. Budburst date was modeled as a quadratic term to test for evidence of stabilizing selection.

|  | A. **RGR 2014-2025, Placerville** | | | B. **RGR 2014-2025, Chico** | | |
| --- | --- | --- | --- | --- | --- | --- |
| *Predictors* | *Estimates* | *CI* | *p* | *Estimates* | *CI* | *p* |
| (Intercept) | 0.26 | 0.22 – 0.30 | **<0.001** | 0.29 | 0.29 – 0.29 | **<0.001** |
| 2014 height | -0.00 | -0.00 – -0.00 | **<0.001** | -0.00 | -0.00 – -0.00 | **<0.001** |
| mean(Budburst) | -0.00 | -0.00 – 0.00 | 0.438 |  |  |  |
| Climate PC1 | 0.00 | 0.00 – 0.00 | **<0.001** | 0.00 | -0.00 – 0.00 | 0.196 |
| mean(Budburst) [1^st^ degree] |  |  |  | -0.01 | -0.10 – 0.07 | 0.754 |
| mean(Budburst) [2^nd^ degree] |  |  |  | -0.11 | -0.19 – -0.03 | **0.008** |
| Observations | 1644 | | | 1708 | | |
| R^2^ / R^2^ adjusted | 0.361 / 0.360 | | | 0.548 / 0.547 | | |

**Table S12**: **Summary of linear mixed model testing the effects of** **budburst date on cumulative 2014-2025 relative growth rates** in *Q. lobata* trees growing in two common gardens. Budburst date was modeled as a quadratic term to test for evidence of stabilizing selection. Climate PC1 and tree height in 2014 were included as covariates to account for known differences in RGR caused by origin climate and tree size prior to outplanting (Goetz *et al.* 2026). Both covariates were scaled to improve comparability. The relationship was tested separately in each garden site; **A**: Placerville garden, **B**: Chico garden.

|  | A. **RGR 2014-2025, Placerville** | | | B. **RGR 2014-2025, Chico** | | |
| --- | --- | --- | --- | --- | --- | --- |
| *Predictors* | *Estimates* | *CI* | *p* | *Estimates* | *CI* | *p* |
| (Intercept) | 0.31 | 0.27 – 0.35 | **<0.001** | 0.35 | 0.31 – 0.39 | **<0.001** |
| 2014 height | -0.00 | -0.00 – -0.00 | **<0.001** | -0.00 | -0.00 – -0.00 | **<0.001** |
| Mean(Leaf elongation) | -0.00 | -0.00 – -0.00 | **0.003** | -0.00 | -0.00 – -0.00 | **0.002** |
| Climate PC1 | 0.00 | 0.00 – 0.00 | **0.001** | 0.00 | -0.00 – 0.00 | 0.461 |
| Observations | 1644 | | | 1708 | | |
| R^2^ / R^2^ adjusted | 0.364 / 0.363 | | | 0.549 / 0.548 | | |

**Table S13: Summary of linear mixed model testing the** **effects of plasticity in budburst date in *Q. lobata*** (measured as coefficient of variation of each tree’s average budburst date over the study period) on cumulative 2014-2025 relative growth rates. Plasticity was modeled as a linear term to test for evidence of plasticity imposing a cost on fitness. Climate PC1 and tree height in 2014 were included as covariates to account for known differences in RGR caused by origin climate and tree size prior to outplanting (Goetz *et al.* 2026). Both covariates were scaled to improve comparability. The relationship was tested separately in each garden site; **A**: Placerville garden, **B**: Chico garden.

|  | A. **RGR 2014-2025, Placerville** | | | B. **RGR 2014-2025, Chico** | | |
| --- | --- | --- | --- | --- | --- | --- |
| *Predictors* | *Estimates* | *CI* | *p* | *Estimates* | *CI* | *p* |
| (Intercept) | 0.19 | 0.17 – 0.21 | **<0.001** | 0.21 | 0.20 – 0.22 | **<0.001** |
| cv(Budburst) | -0.13 | -0.27 – 0.02 | 0.081 | -0.02 | -0.08 – 0.03 | 0.401 |
| Climate PC1 | 0.01 | 0.00 – 0.01 | **<0.001** | 0.00 | -0.00 – 0.00 | 0.457 |
| 2014 height | -0.04 | -0.04 – -0.04 | **<0.001** | -0.05 | -0.05 – -0.04 | **<0.001** |
| Observations | 623 | | | 626 | | |
| R^2^ / R^2^ adjusted | 0.372 / 0.369 | | | 0.565 / 0.563 | | |

**Table S14: Summary of linear mixed model testing the effects of plasticity in leaf elongation date** (measured as coefficient of variation of each tree’s average leaf elongation date over the study period) **on cumulative 2014-2025 relative growth rates**. Plasticity was modeled as a linear term to test for evidence of plasticity imposing a cost on fitness. The relationship was tested separately in each garden site; **A**: Placerville garden, **B**: Chico garden.

|  | A. **RGR 2014-2025, Placerville** | | | B. **RGR 2014-2025, Chico** | | |
| --- | --- | --- | --- | --- | --- | --- |
| *Predictors* | *Estimates* | *CI* | *p* | *Estimates* | *CI* | *p* |
| (Intercept) | 0.17 | 0.15 – 0.19 | **<0.001** | 0.20 | 0.19 – 0.21 | **<0.001** |
| cv(Leaf elongation) | 0.00 | -0.12 – 0.12 | 0.991 | 0.05 | -0.07 – 0.18 | 0.386 |
| Climate PC1 | 0.01 | 0.00 – 0.01 | **<0.001** | 0.00 | -0.00 – 0.00 | 0.502 |
| 2014 height | -0.04 | -0.04 – -0.03 | **<0.001** | -0.05 | -0.05 – -0.04 | **<0.001** |
| Observations | 623 | | | 626 | | |
| R^2^ / R^2^ adjusted | 0.369 / 0.366 | | | 0.565 / 0.563 | | |
